# Low-self-reactive, circulation-biased blood regulatory T cells sense danger-associated nucleotides to restrain atherosclerosis

**DOI:** 10.64898/2026.05.07.723421

**Authors:** Takashi Sekiya, Shinya Hidano, Satoshi Takaki

## Abstract

Peripheral blood (PB) is the predominant source of regulatory T cells (Tregs) for clinical applications, yet PB Tregs have not been systematically compared with tissue Tregs and their defining properties remain poorly understood. Here we identify a circulation-biased PB Treg population with reduced self-reactivity. Although PB Tregs are less suppressive in vitro, they are hypersensitive to the danger-associated nucleotides ATP and NAD^+^, linked to a high ARTC2.2/CD38 ratio. ATP/NAD^+^-triggered apoptotic conversion of PB Tregs promotes macrophage efferocytosis and selectively dampens IFN-γ–driven inflammatory polarization while preserving IL-10–associated programs and enhancing the scavenger receptor Mertk, suggesting that efferocytosis-mediated macrophage reprogramming contributes to lesion control. Consistent with this model, in atherosclerosis-prone mice lacking T cells, PB Treg transfer effectively limits plaque growth and necrotic core formation. We identify an ATP/NAD^+^-hypersensitive PB Treg fraction in both mice and humans and provide evidence consistent with preferential efferocytosis of this subset within atherosclerotic plaques.

## Introduction

Regulatory T cells (Tregs) enforce immune tolerance and are increasingly pursued as cell therapies because they can impose more antigen-selective immunosuppression than conventional immunosuppressive drugs.(Sakaguchi *et al*, 2020; Tang & Bluestone, 2013) In most clinical settings, peripheral blood (PB) is the dominant source of Tregs—either used directly or after ex vivo expansion—because it is readily accessible. Despite this translational importance, PB Tregs have rarely been subjected to the kind of systematic, cross-tissue comparisons that have illuminated specialized Treg programs in various other sites including adipose tissue, skeletal muscle, and the central nervous system.(Burzyn *et al*, 2013; Feuerer *et al*, 2009; Ito *et al*, 2019) In humans, matched sampling of tissue Tregs at high quality is difficult; in mice, blood Tregs have often been treated simply as cells in transit. Consequently, the defining properties and physiological roles of blood-resident Tregs remain incompletely understood.

Tregs are distinguished from conventional CD4 T cells (Tconv) in multiple ways. In the thymus, relatively strong recognition of self-antigen favors commitment to the Foxp3^+^ lineage, enriching the peripheral Treg pool for self-reactive TCRs.(Hsieh *et al*, 2006; Jordan *et al*, 2001; Josefowicz *et al*, 2012; Sekiya *et al*, 2013) Consistent with this concept, peripheral Tregs typically show higher tonic TCR signaling than Tconv cells, a feature that contributes to immune homeostasis and autoimmunity prevention.(Levine *et al*, 2014; Moran *et al*, 2011) Another reported hallmark—particularly in mice—is heightened sensitivity to danger-associated nucleotides, most notably ATP and NAD^+^, which can provoke P2X7R-dependent apoptotic death of Tregs.(Aswad *et al*, 2005; Hubert *et al*, 2010) This vulnerability has been exploited experimentally to transiently deplete Tregs and enhance antitumor immunity, but its physiological or disease-relevant meaning remains unclear.(Hubert *et al*., 2010)

Tregs also constrain atherosclerosis, an inflammatory disease driven by both innate and adaptive immune responses.(Hansson & Hermansson, 2011; Witztum & Lichtman, 2014) Macrophages play a particularly central role in atherogenesis. IFN-γ–polarized inflammatory macrophages promote plaque growth and destabilization by producing pro-inflammatory cytokines and chemokines (e.g., TNF-α, IL-1β, and MCP-1/CCL2) and by inducing tissue-injurious mediators such as matrix metalloproteinases and iNOS, whereas macrophages polarized by IL-10, TGF-β, or IL-13 are thought to exert reparative and anti-inflammatory functions through expression of scavenger receptors (e.g., CD163, CD206, and Mertk) and anti-inflammatory mediators such as IL-10 and TGF-β, thereby contributing to plaque stabilization and limiting lesion progression.(Chinetti-Gbaguidi *et al*, 2015; Moore *et al*, 2013; Ouyang & Liu, 2024) In mouse models, Treg depletion accelerates plaque progression, whereas adoptive transfer of Tregs is broadly protective.(Ait-Oufella *et al*, 2006; Klingenberg *et al*, 2013; Mor *et al*, 2007; Sharma *et al*, 2020) Proposed mechanisms include IL-10– and TGF-β–dependent modulation of macrophage polarization, IL-13–mediated enhancement of efferocytosis of dying cells, and suppression of pro-atherogenic T cell responses, including Th1 differentiation and reactivity to specific antigens such as ApoB100-derived antigens.(Herbin *et al*, 2012; Mallat *et al*, 2007; Ouyang & Liu, 2024; Proto *et al*, 2018) In addition, Treg-derived exosomes have been reported to act directly on macrophages, suppressing expression of M1-associated markers such as TNF-α and iNOS while inducing M2-associated markers including Arg1 and TGF-β, thereby conferring protection in murine models of acute myocardial infarction. (Hu *et al*, 2020) These observations suggest that Tregs contain molecular constituents capable of directly reprogramming macrophage inflammatory states. However, the complete mechanistic repertoire of atheroprotective Tregs is not yet fully defined. Of note, multiple clinical studies have reported a reduction in circulating FOXP3^+^ Tregs in patients with atherosclerotic cardiovascular disease.(Wigren *et al*, 2012; Yazdani *et al*, 2016) Comparable decreases in Treg abundance have also been observed during atherogenesis in ApoE- and LDLR-deficient mice.(Maganto-Garcia *et al*, 2011; Mor *et al*., 2007) However, it remains unclear whether a discrete Treg subset is preferentially diminished in the context of atherosclerosis and, if so, what the biological consequences of such selective loss are.

Here we focus on PB Tregs. We identify a circulation-biased Treg population that is enriched for low self-reactivity and exhibits attenuated suppressive function, yet displays pronounced nucleotide hypersensitivity. We further show that ATP/NAD^+^-triggered apoptotic PB Tregs are efferocytosed by macrophages, which in turn selectively dampen inflammatory polarization. In an atherosclerosis model, PB Tregs traffic efficiently to plaques and suppress plaque development via an ARTC2.2-dependent program, supporting nucleotide-driven Treg apoptosis and macrophage efferocytosis as a previously underappreciated immunoregulatory axis.

## Results

### PB Tregs are enriched for low ongoing TCR signaling and low self-reactivity

We first surveyed eight tissues and confirmed that Foxp3^+^ Tregs can be readily detected in PB, although they comprise a smaller fraction of CD4^+^ T cells than in most tissues (**Supplementary Fig. S1A, S1B**). We then performed ATAC-seq on naïve CD4 T cells and Tregs from six tissues, including blood. Principal component analysis revealed that PB Tregs segregate from other tissue Tregs and occupy an intermediate position between naïve T cells and Tregs along PC1 (**Fig. 1A**), suggesting a distinct epigenomic state.

**Figure 1.**
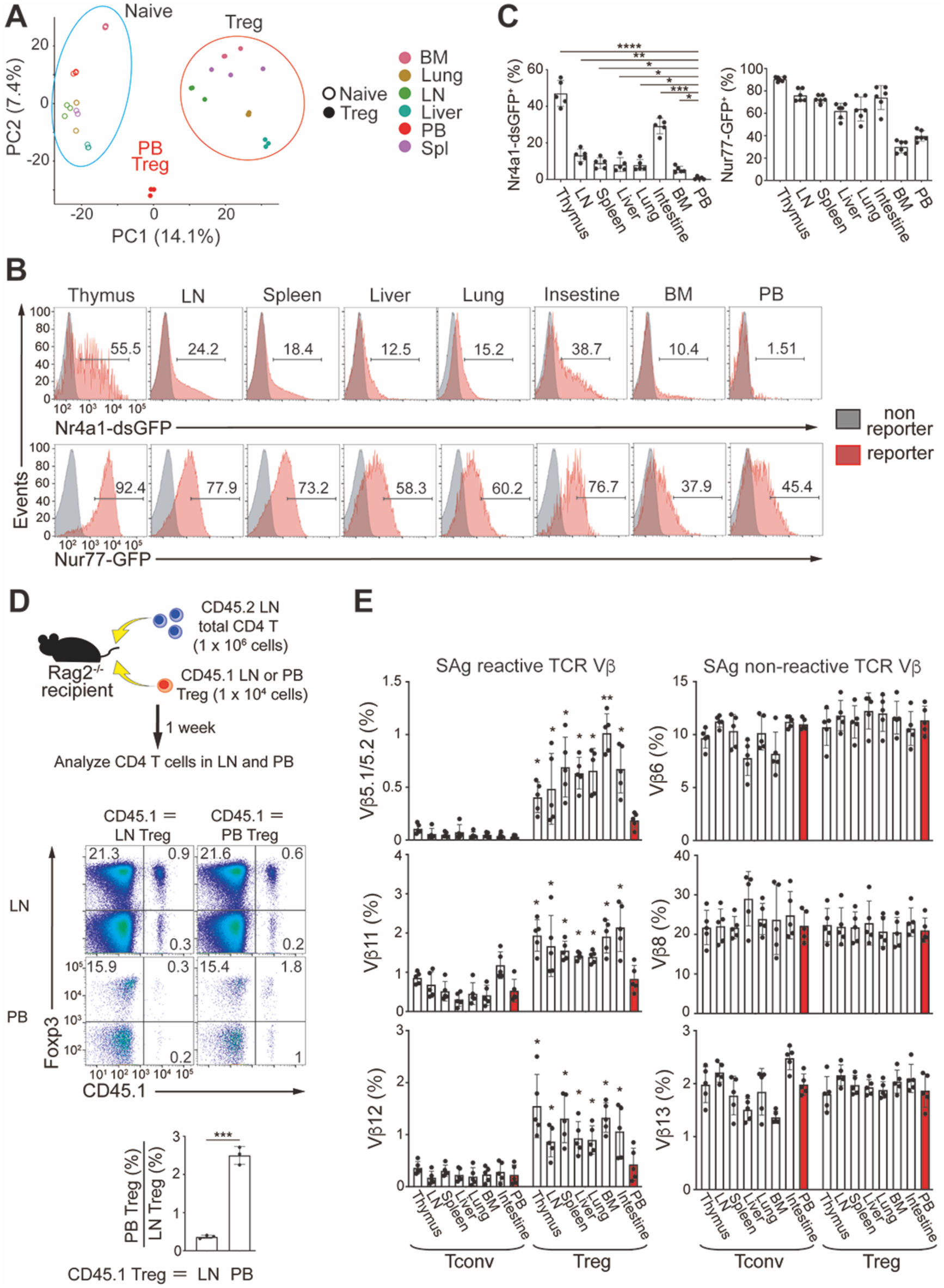
Peripheral blood Tregs are composed of circulation-biased cells and enriched for cells with reduced self-reactivity. **(A)** Principal component analysis of ATAC-seq profiles from naïve CD4 T cells and Tregs isolated from the indicated organs. BM: bone marrow; LN: lymph nodes; PB: peripheral blood; Spl: spleen. **(B)** Representative histograms of Nr4a1-dsGFP (destabilized reporter) and conventional Nur77-GFP (stable EGFP reporter) signals in Tregs across tissues. Numbers adjacent to outlined areas indicate percent GFP^+^ cells. **(C)** Quantification of Nr4a1-dsGFP^+^ and Nur77-GFP^+^ Treg frequencies across tissues. Each dot represents an individual mouse, mean ± SD (n = 6). **(D)** Adoptive transfer into Rag2^−/−^ recipients of total CD4 T cells (CD45.2) plus purified Tregs (CD45.1) derived from LN or PB, followed by quantification of CD45.1 Treg enrichment in PB versus LN at 1 week. Top: A schematic of experiments shown in this panel. Middle: Flow cytometry of total CD4^+^ T cells, obtained from LN or PB of the mice transferred with the indicated combination of the donor cells. Bottom: Quantification of the results. Data are representative of three biological replicates performed in three independent experiments, set up in triplicate, mean ± SD. **(E)** Frequencies of superantigen-reactive versus non-reactive TCR Vβ subsets among Tconv and Treg populations across tissues (BALB/c). Each dot represents an individual mouse, mean ± SD (n = 5). \**p* < 0.05; ** *p* < 0.01; *** *p* < 0.005; **** *p* < 0.001, one-way ANOVA with the Bonferroni test **(C)**, **(E)**, Student’s t test **(D)**.

Because Tregs are typically sustained by ongoing self-antigen recognition, we next compared tonic T cell receptor (TCR) signaling across tissues. We used a recently generated Nr4a1 (Nur77) reporter mouse (Nr4a1-dsGFP mice) in which a destabilized GFP (half-life of a few hours) reports recent TCR signaling, enabling detection of ongoing stimulation in situ.(Sekiya *et al*, 2024) Strikingly, the fraction of Nr4a1-dsGFP^+^ Tregs was markedly lower in blood than in other tissues (**Fig. 1B, 1C**). In contrast, in conventional Nur77 reporter mice (Nur77-GFP mice), in which EGFP is used as a reporter molecule and is relatively stable (half-life >24 h),(Moran *et al*., 2011) a non-trivial fraction of PB Tregs remained GFP^+^ (**Fig. 1B, 1C**). These results collectively suggest that PB Tregs are composed of a mixture of long-lived blood-biased Tregs and recently emigrated tissue Tregs.

To test whether PB Tregs merely recirculate or preferentially persist in blood, we adoptively transferred total CD4 T cells (CD45.2, 1×10^6^) together with a small number of purified Tregs (CD45.1, 1×10^4^) from either lymph nodes or blood into Rag2^−/−^ recipients. One week later, PB-derived Tregs showed significantly greater enrichment in the circulation relative to LN-derived Tregs (**Fig. 1D**), supporting the presence of a circulation-biased Treg population.

We next assessed self-reactivity of PB Tregs at the TCR level employing BALB/c mice that express endogenous superantigens encoded by Mtv-8 and Mtv-9, that react with specific subsets of Vβ chains (e.g., Vβ5, Vβ11, and Vβ12). Superantigen-reactive Vβ usage is known to be typically enriched among Tregs relative to Tconv cells.(Tanaka *et al*, 2010) In this experiment, superantigen-reactive Vβ usage has been estimated as a surrogate for heightened self-antigen recognition. Using the Nr4a1-dsGFP mice, it was validated that tissue Tregs with superantigen-reactive Vβ subsets indeed display stronger tonic TCR signaling (**Supplementary Fig. S1C, S1D**). As shown in **Fig. 1E**, we found that superantigen-reactive Vβ subsets were under-represented among PB Tregs, whereas non-reactive Vβ subsets were comparable across tissues. This tissue bias was not observed in C57BL/6J mice, which lack I-E and thus cannot present the relevant superantigens (**Supplementary Fig. S1E**).

To functionally test self-reactivity, we performed autologous mixed lymphocyte reactions (MLR) using LN- or PB-derived Tregs from Nr4a1-dsGFP mice. While polyclonal anti-CD3/anti-CD28 stimulation elicited comparable activation, PB Tregs showed markedly reduced responses in autologous MLR (**Supplementary Fig. S1F, S1G**). Together, these data indicate that the PB Treg compartment is enriched for Tregs with low ongoing TCR stimulation and reduced self-reactivity.

We next performed RNA-seq on Tregs from six tissues including blood. Tregs clustered by tissue of origin, indicating distinct gene expression programs (**Supplementary Fig. S2A, S2B**). Gene Ontology (GO) analysis of the PB-specific gene cluster (Cluster 3) highlighted terms such as “platelet activation,” “blood coagulation,” and “hemostasis” (**Supplementary Fig. S2C**)), suggesting that PB Tregs may be shaped by, and potentially tuned to, the circulating milieu. Genes in each cluster used for GO analysis are listed in **Supplementary Table S1**. Another PB-enriched gene cluster (Cluster 1) was enriched with tissue-tropic factors like chemokine receptors, and GO analysis of this cluster highlighted terms such as “leukocyte migration” and “leukocyte chemotaxis”, implicating their tropism to inflammatory sites.

### PB Tregs have heightened sensitivity to danger-associated nucleotides ATP and NAD^+^

During assay development we noticed that PB Tregs differed dramatically depending on how red blood cells (RBC) were removed. Compared with density gradient separation, hypotonic RBC lysis induced robust apoptosis in PB Tregs, as indicated by phosphatidylserine exposure (**Fig. 2A, 2B**). This effect did not seem to be caused by an artificial toxicity of RBC lysis, because PB Tconv cells as well as the majority of cells in PB were far more resistant to the treatment (**Fig. 2A-2D**). In addition, RBC removal from PB was performed by density-gradient separation throughout this study, unless otherwise indicated.

**Figure 2.**
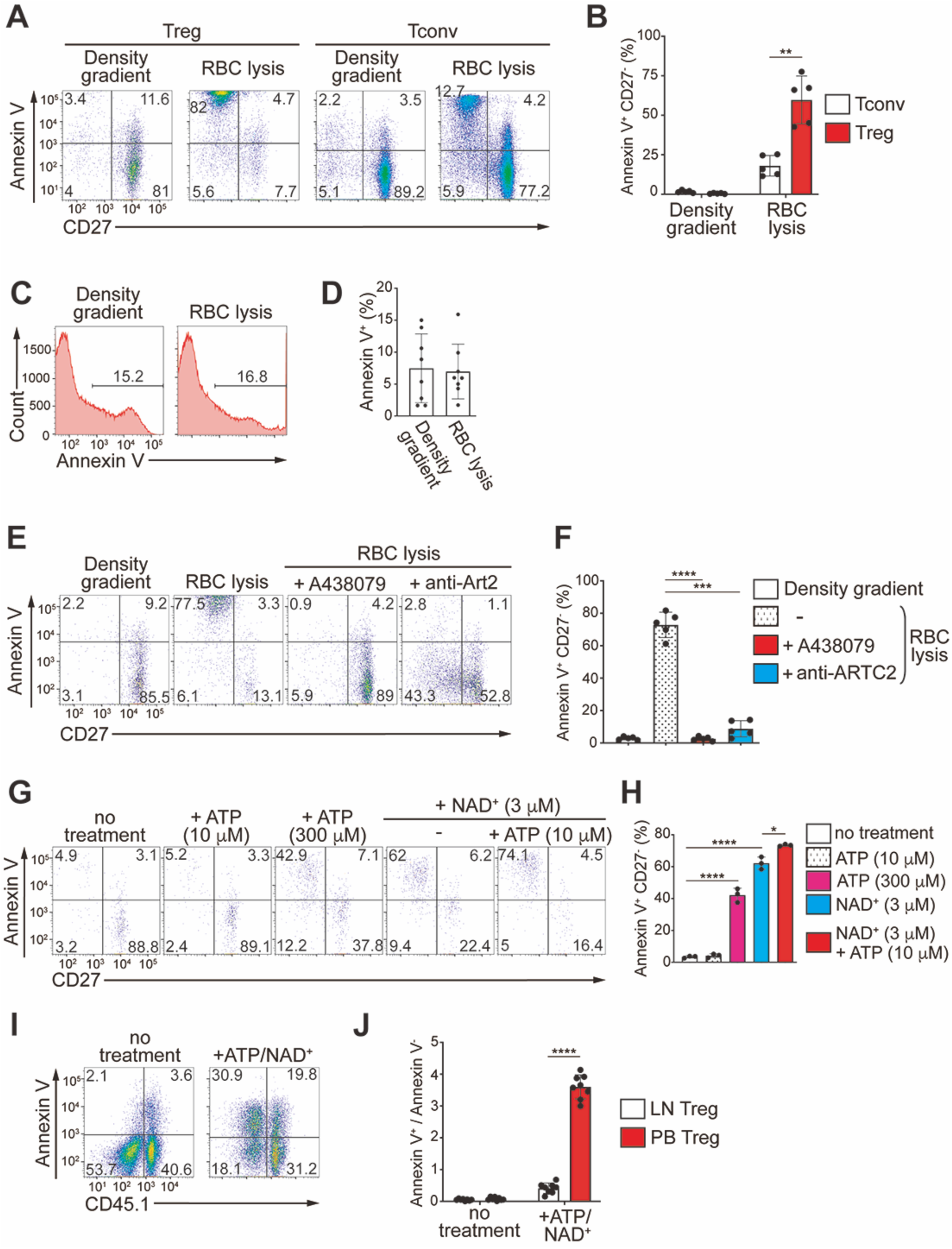
PB Tregs have heightened sensitivity to danger-associated nucleotides ATP/NAD^+^: **(A)** Flow cytometry of Annexin V versus CD27 on peripheral blood (PB) Tregs and PB Tconv cells after red blood cell (RBC) removal by density gradient or hypotonic lysis. Numbers indicate the percentage of cells in the indicated regions. **(B)** Quantification of the results shown in **(A)**. Each dot represents an individual mouse, mean ± SD (n = 5). **(C, D)** Flow cytometry of Annexin V expression **(C)** and quantification **(D)** of total PB cells after removal of RBC by density gradient vs hypotonic lysis. Each dot represents an individual mouse, mean ± SD (n = 8). **(E, F)** Flow cytometry of PB Tregs after removal of RBC by density gradient or hypotonic lysis performed with P2X7R inhibitor (A438079) or anti-ARTC2.2 blockade; representative plots **(E)** and quantification **(F)**. Data were pooled from five independent biological replicates, performed in three independent experiments, mean ± SD. **(G, H)** Flow cytometry of in vitro ATP and/or NAD^+^-treated PB Tregs **(G)** and quantification of Annexin V^+^ CD27^−^ cells **(H)**. Data are representative of three biological replicates performed in three independent experiments, set up in triplicate, mean ± SD. **(I**, **J)** Paired comparison of PB (CD45.2) versus LN (CD45.1) Tregs treated with ATP/NAD^+^ in a same reaction; representative plots **(I)** and quantification **(J)**. Data were pooled from eight independent biological replicates performed in four independent experiments, mean ± SD. \**p* < 0.05; ** *p* < 0.01; *** *p* < 0.005; **** *p* < 0.001, one-way ANOVA with the Bonferroni test **(B)**, **(F)**, **(H)**, **(J).**

Next, to elucidate the mechanism underlying the Treg-selective induction of apoptosis by RBC lysis procedure, we focused on the known property that Tregs are more sensitive than Tconv cells to stimulation by the danger-associated molecular patterns (DAMPs) nucleotides ATP and NAD^+^.(Aswad *et al*., 2005; Hubert *et al*., 2010) Both ATP and NAD^+^ are released predominantly from dying cells in inflamed tissues and function as danger-associated signals. We therefore performed the RBC lysis in the presence of the P2X7R inhibitor A438079, which blocks ATP-driven signaling, and a neutralizing antibody against ARTC2.2, a key mediator of NAD^+^-dependent signaling. Under these conditions, both reagents markedly suppressed apoptosis in Tregs, indicating that the hemolysis-induced apoptosis of Tregs is driven by ATP and/or NAD^+^ released from RBCs (**Fig. 2E, 2F**).

To define the relative contribution of ATP and NAD^+^ to Treg apoptosis, we next performed in vitro stimulation assays using Tregs prepared by density gradient separation. Based on reports that extracellular NAD^+^ concentrations can rise to approximately 1–10 µM in inflamed tissues,(Adriouch *et al*, 2007) we used 3 µM NAD^+^, and tested ATP at 10 µM and 300 µM (10 minutes, room temperature). As reported previously, NAD^+^ alone robustly induced apoptosis at 3 µM (**Fig. 2G, 2H**).(Aswad *et al*., 2005) In contrast, ATP alone had no detectable effect at 10 µM, and even at 300 µM induced weaker apoptosis than 3 µM NAD^+^, consistent with prior observations (**Fig. 2G, 2H**).(Aswad *et al*., 2005) Importantly, ATP exhibited the expected cooperativity with NAD^+^:(Seman *et al*, 2003) 10 µM ATP potentiated the apoptotic effect of 3 µM NAD^+^ (**Fig. 2G, 2H**). Based on these results, subsequent experiments were performed using 10 µM ATP together with 3 µM NAD^+^ (hereafter referred to as ATP/NAD^+^).

Finally, we compared PB and LN Tregs in an identical in vitro nucleotide challenge. Under matched conditions, PB Tregs were significantly more sensitive to ATP/NAD^+^ than LN Tregs (**Fig. 2I, 2J**), indicating that heightened nucleotide sensitivity is an intrinsic feature of PB Tregs.

### Low tonic TCR signaling associates with a high ARTC2.2/CD38 ratio and heightened nucleotide sensitivity of PB Tregs

Next, we investigated the mechanism underlying the heightened sensitivity of PB Tregs to ATP/NAD^+^. Given our observation that PB Tregs experience weaker self-antigen–driven stimulation, we asked whether tonic TCR signaling modulates ATP/NAD^+^ responsiveness. To this end, LN Tregs from Nur77-GFP reporter mice were stratified into four equal fractions based on GFP intensity and challenged with ATP/NAD^+^. Strikingly, GFP intensity inversely correlated with ATP/NAD^+^ sensitivity (**Fig. 3A–3C**), strongly suggesting that tonic TCR signaling dampens nucleotide-induced susceptibility in Tregs. In addition, Nur77-GFP intensity in Tregs was not altered by ATP/NAD^+^ treatment (**Fig. 3D**), showing that the reporter signal reflects antecedent TCR stimulation rather than an acute response to nucleotide exposure.

**Figure 3.**
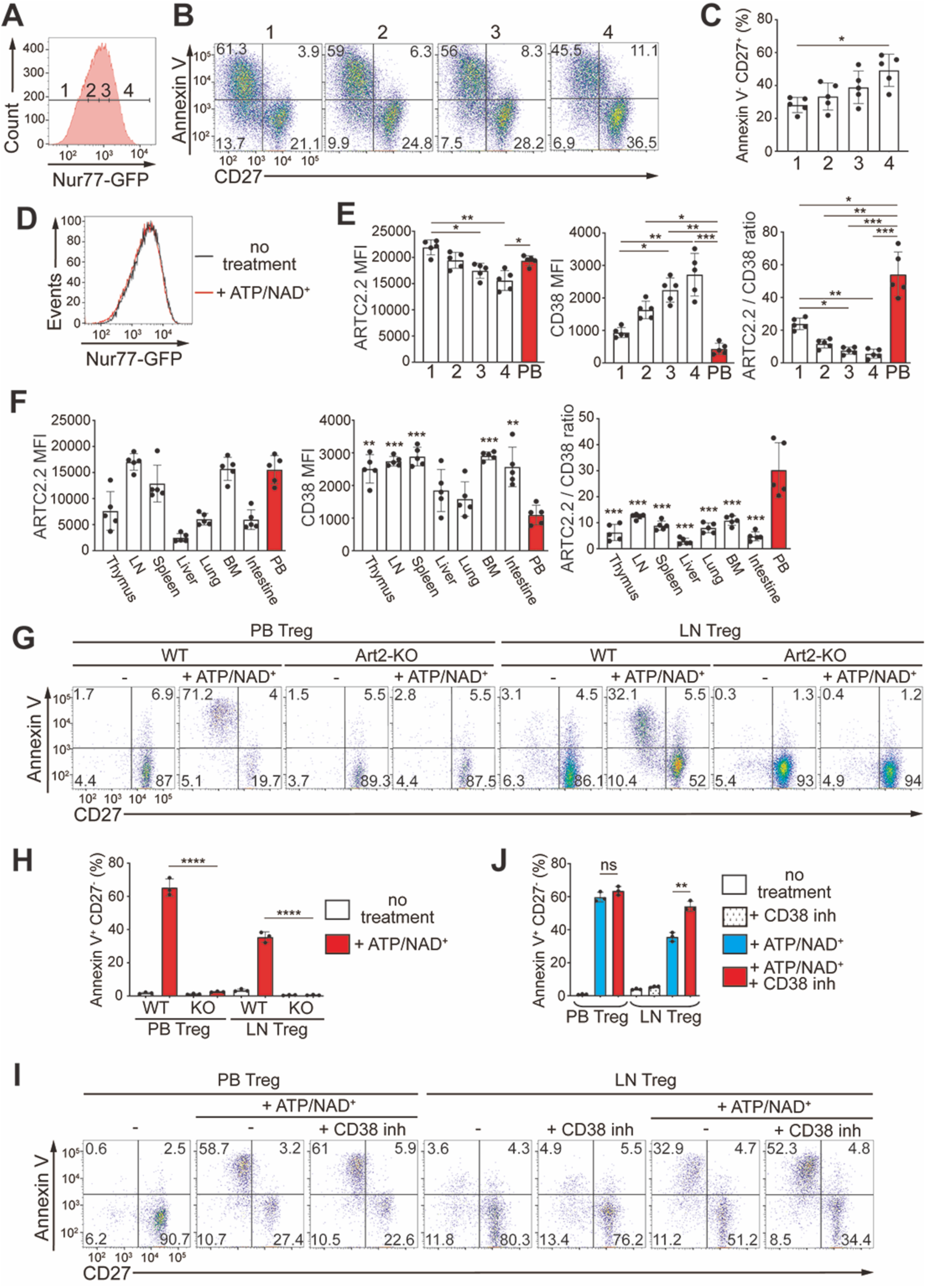
Low tonic TCR signaling associates with a high ARTC2.2/CD38 ratio and heightened nucleotide sensitivity of PB Tregs: **(A)** Nur77-GFP intensity–based stratification of lymph nodes (LN) Tregs into four fractions. **(B, C)** Flow cytometry of Annexin V versus CD27 after ATP/NAD^+^ challenge of LN Tregs in each fraction **(B)** and quantification showing an inverse correlation between Nur77-GFP intensity and nucleotide sensitivity **(c)**. Numbers indicate the percentage of cells in the indicated regions **(B)**. Data were pooled from five independent biological replicates, mean ± SD. **(D)** Flow cytometry histogram showing that Nur77-GFP intensity of Tregs is unchanged by ATP/NAD^+^ treatment. **(E)** ARTC2.2 mean fluorescence intensity (MFI), CD38 MFI, and ARTC2.2/CD38 ratio determined by flow cytometry analysis across Nur77-GFP fractions of LN Tregs and PB Tregs. Each dot represents an individual mouse, mean ± SD (n = 5). **(F)** ARTC2.2 MFI, CD38 MFI, and ARTC2.2/CD38 ratio of Tregs determined by flow cytometry analysis across organs. Each dot represents an individual mouse, mean ± SD (n = 5). **(G, H)** Flow cytometry analysis of ATP/NAD^+^-induced apoptotic conversion in WT versus Art2-KO PB and LN Tregs **(G)** and quantification **(H)**. Data are representative of three biological replicates performed in three independent experiments, set up in triplicate, mean ± SD. WT: wild-type. **(I, J)** Flow cytometry analysis of effect of CD38 inhibition on ATP/NAD^+^-induced apoptosis in PB and LN Tregs **(I)** and quantification **(J)**. Data are representative of three biological replicates performed in three independent experiments, set up in triplicate, mean ± SD. \**p* < 0.05; ** *p* < 0.01; *** *p* < 0.005; **** *p* < 0.001, one-way ANOVA with the Bonferroni test **(C)**, **(E)**, **(F)**, **(H)**, **(J).**

Given the inverse correlation between tonic TCR signal strength and ATP/NAD^+^ sensitivity, we next sought to define the underlying molecular mechanism. Because NAD^+^ is the dominant trigger of Treg apoptosis, we quantified expression of ARTC2.2—an NAD^+^-dependent ecto-ADP-ribosyltransferase that functions as the key receptor/effector upstream of P2X7R activation—and CD38, an ectoenzyme that degrades extracellular NAD^+^.(Rivas-Yanez *et al*, 2020) We analyzed these molecules across the four Nur77-GFP–stratified LN Treg fractions described above and compared them with PB Tregs. ARTC2.2 expression inversely correlated with Nur77-GFP intensity, whereas CD38 expression showed a positive correlation (**Fig. 3E**). Notably, PB Tregs exhibited a markedly higher ARTC2.2/CD38 ratio than LN Tregs, providing a mechanistic basis for their enhanced nucleotide sensitivity. Furthermore, PB Tregs exhibited the highest ARTC2.2/CD38 ratio among tissues (**Fig. 3F**).

Genetic deletion of *Art2* (both ARTC2.2 and Art2a) abrogated ATP/NAD^+^-induced apoptosis in both PB and LN Tregs, as reported previously (**Fig. 3G, 3H**).(Hubert *et al*., 2010) Conversely, pharmacologic inhibition of CD38 selectively increased nucleotide sensitivity in LN Tregs but had little effect on PB Tregs (**Fig. 3I, 3J**), consistent with the already-low CD38 state in blood. These findings suggest that weak tonic TCR signaling contributes to the heightened sensitivity of PB Tregs by skewing the ARTC2.2/CD38 axis.

### PB Tregs have reduced suppressive capacity in vitro

Next, we compared the in vitro suppressive capacity of PB Tregs and LN Tregs in standard T cell proliferation–suppression assays. PB Tregs suppressed responder T cell proliferation significantly less effectively than LN Tregs (**Supplementary Fig. 3A, 3B**). To investigate the basis for this functional difference, we leveraged our observation that PB Tregs receive weaker self-antigen–driven stimulation and tested whether tonic TCR signaling influences suppressive potency. LN Tregs from Nur77-GFP reporter mice were stratified into four equal fractions according to GFP intensity, and their suppressive activity was assessed in parallel (**Supplementary Fig. S3C**). Suppressive capacity positively correlated with Nur77-GFP intensity, strongly suggesting that tonic TCR signaling enhances the ability of Tregs to suppress T cell proliferation (**Supplementary Fig. S3D, S3E**). To gain mechanistic insight, we quantified expression of key mediators implicated in in vitro Treg suppression—CD25, CTLA-4, and IL-10—in the Nur77-GFP–stratified LN Treg fractions and in PB Tregs. CD25 and IL-10 expression positively correlated with Nur77-GFP intensity, and both were reduced in PB Tregs (**Supplementary Fig. S3F, S3G**). CTLA-4 expression did not correlate with Nur77-GFP intensity across LN fractions, but was nonetheless lower in PB Tregs than in LN Tregs (**Supplementary Fig. S3F**). Collectively, these findings suggest that the weak tonic TCR stimulation experienced by PB Tregs is associated with reduced expression of CD25 and IL-10 (and lower CTLA-4 overall), which likely contributes to their weakened capacity to suppress T cell proliferation in vitro.

### Efferocytosis of apoptotic PB Tregs selectively suppresses inflammatory macrophage polarization

The apparent functional deficit of PB Tregs raised the possibility that blood is merely a reservoir of poorly functional Tregs. We therefore asked whether PB Tregs might instead exert distinct, non-canonical immunoregulatory functions. Because apoptotic cells are efficiently engulfed by macrophages and can reprogram their activation states,(Elliott & Ravichandran, 2016) we hypothesized that nucleotide-triggered apoptotic PB Tregs could modulate macrophage polarization via efferocytosis.

To isolate macrophages that had engulfed specific T cell populations, previous work has typically relied on amine-reactive cytosolic dyes; however, because these dyes such as CFSE covalently modify intracellular proteins and can potentially perturb their functions, we screened fluorochrome-conjugated anti-CD4 antibodies for durable retention after phagocytosis (**Supplementary Fig. S4A**). Only Brilliant Violet (BV) 421 and Alexa Fluor 647 signals persisted within bone marrow-derived macrophages (BMDMs) 48 h after engulfment (**Supplementary Fig. S4B**). Uptake of ATP/NAD^+^-treated CD27^−^ Tregs was inhibited by recombinant Annexin V (**Supplementary Fig. S4C, S4D**). While antibody labeling may provide opsonizing signals that facilitate uptake, this Annexin V sensitivity shows that phosphatidylserine-dependent efferocytosis did contribute to this process. Calcein-AM assays indicated that ATP/NAD^+^-treated Tregs and Tconv cells were still largely viable at the time of mixing with BMDMs, despite Annexin V positivity (**Supplementary Fig. S4E, S4F**), consistent with early apoptotic conversion.

Based on the above preparative considerations, we established a tri-culture system in which BMDMs were co-incubated with BV421- or Alexa647-labeled Tregs and Alexa647 or BV421-labeled Tconv cells (both ATP/NAD^+^-treated CD27^−^ cells), followed by polarization with IFN-γ (M1-like) or IL-10 (M2c-like), and sorting of phagocytic versus non-phagocytic macrophages (**Fig. 4A, 4B**). Transcriptomic profiling showed that Treg cell-phagocytic macrophages separated clearly from non-phagocytic, Tconv cell-phagocytic, and baseline (M0) macrophages (**Fig. 4C, 4D**). Notably, Euclidean distance analysis across samples revealed that Treg-phagocytic macrophages were closer to the M0 state than were other IFN-γ–polarized conditions (**Fig. 4C**), consistent with the notion that Treg engulfment negatively regulates IFN-γ–driven polarization. In IFN-γ–polarized macrophages, we analyzed the expression of inflammatory M1 marker genes and found that engulfment of Tregs suppressed induction of multiple transcripts, including *Nos2* (iNOS) (**Fig. 4E**). In IL-10–polarized macrophages we examined expression of M2c-associated genes implicated in immunosuppression and resolution of inflammation. Although we did not observe Treg-phagocytosis–specific changes in this program, increased expression of efferocytosis-related genes such as Mertk was detected in both Treg-engulfing and Tconv-engulfing macrophages (**Fig. 4F**). Regulation of *Nos2* and *Mertk* was independently reproduced by qPCR (**Fig. 4G**). For the above RNA-seq experiments, Tregs and Tconv cells were isolated from lymph nodes and spleen to obtain sufficient cell numbers; importantly, PB–derived Tregs and Tconv cells elicited indistinguishable effects, demonstrating that the observed macrophage reprogramming is not specific to lymphoid-tissue–derived T cells.

**Figure 4.**
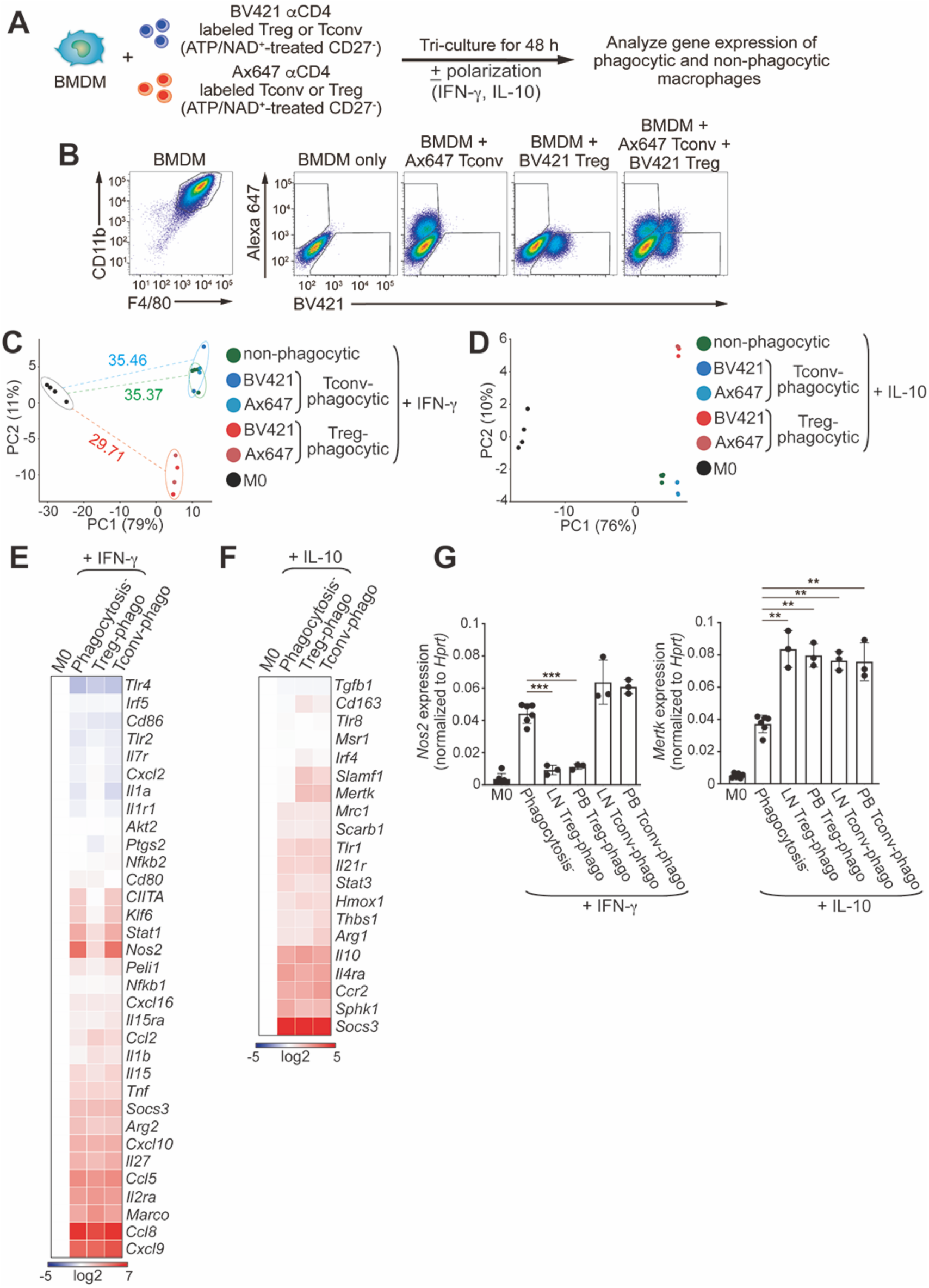
Efferocytosis of apoptotic Tregs selectively suppresses IFN-γ–driven inflammatory macrophage polarization: **(A)** Experimental scheme for tri-culture: BMDMs co-incubated with BV421-or Alexa647-anti-CD4–labeled ATP/NAD^+^-treated CD27^−^ Tregs and Tconv cells, followed by polarization with IFN-γ or IL-10 and sorting of phagocytic versus non-phagocytic macrophages. **(B)** Representative flow cytometry gating to identify macrophages (F4/80^+^CD11b^+^) and phagocytic subsets (BV421^+^/Alexa647^+^). **(C)** Principal component analysis (PCA) of RNA-seq result obtained with IFN-γ polarization. Euclidean distances between non-polarized (M0) and IFN-γ-polarized macrophages, either non-phagocytic (green), Tconv-phagocytic (cyan), or Treg-phagocytic (red) are shown. **(D)** PCA of RNA-seq result obtained with IL-10 polarization. **(E)** Heatmap of RNA-seq result obtained with IFN-γ polarization showing expression of inflammatory M1 macrophage marker genes in the indicated cells. phago: phagocytic. **(F)** Heatmap of RNA-seq result obtained with IL-10 polarization showing expression of anti-inflammatory reparative M2c macrophage marker genes in the indicated cells. **(G)** qPCR analysis of *Nos2* and *Mertk* expression levels, normalized to *Hprt*, in the indicated cells. Data are representative of three biological replicates performed in three independent experiments, set up in triplicate, mean ± SD. ** *p* < 0.01; *** *p* < 0.005, one-way ANOVA with the Bonferroni test.

### Atherosclerosis reduces the nucleotide-hypersensitive PB Treg subset

As described above, PB Tregs exhibit a propensity to preferentially persist within the circulation, raising the possibility that—by virtue of their localization—they may participate in patrolling the vascular endothelium. Tregs are known to exert protective, suppressive functions during atherosclerotic plaque progression; however, many aspects of the underlying mechanisms remain incompletely defined. We therefore investigated PB Tregs in an ApoE-deficient mouse model of atherosclerosis (B6.SHL).(Matsushima *et al*, 1999)

Previous studies have reported that ApoE deficiency is associated with a reduction in Treg abundance, particularly in aged mice, yet it has remained unclear which Treg subpopulation(s) are preferentially lost.(Maganto-Garcia *et al*., 2011; Mor *et al*., 2007) In this study, we first found that PB Treg numbers dropped at approximately 18 weeks of age—an interval during which plaque formation becomes prominent(Reddick *et al*, 1994)—in ApoE-deficient mice (**Fig. 5A**). In parallel with this decline, PB Tregs exhibited increased CD38 mean fluorescence intensity (MFI) and decreased ARTC2.2 MFI around 18 weeks of age (**Fig. 5B, 5C**). We next compared ATP/NAD^+^ sensitivity of PB Tregs from wild-type and ApoE-deficient mice at 6 weeks of age and at the experimental endpoint of 26 weeks. Whereas no difference was evident at 6 weeks, PB Tregs from 26-week-old ApoE-deficient mice showed a marked reduction in sensitivity to ATP/NAD^+^ stimulation (**Fig. 5D, 5E**). Collectively, these findings suggest that, as atherosclerosis progresses in ApoE-deficient mice, the ATP/NAD^+^-hypersensitive fraction of PB Tregs may be selectively eliminated.

**Figure 5.**
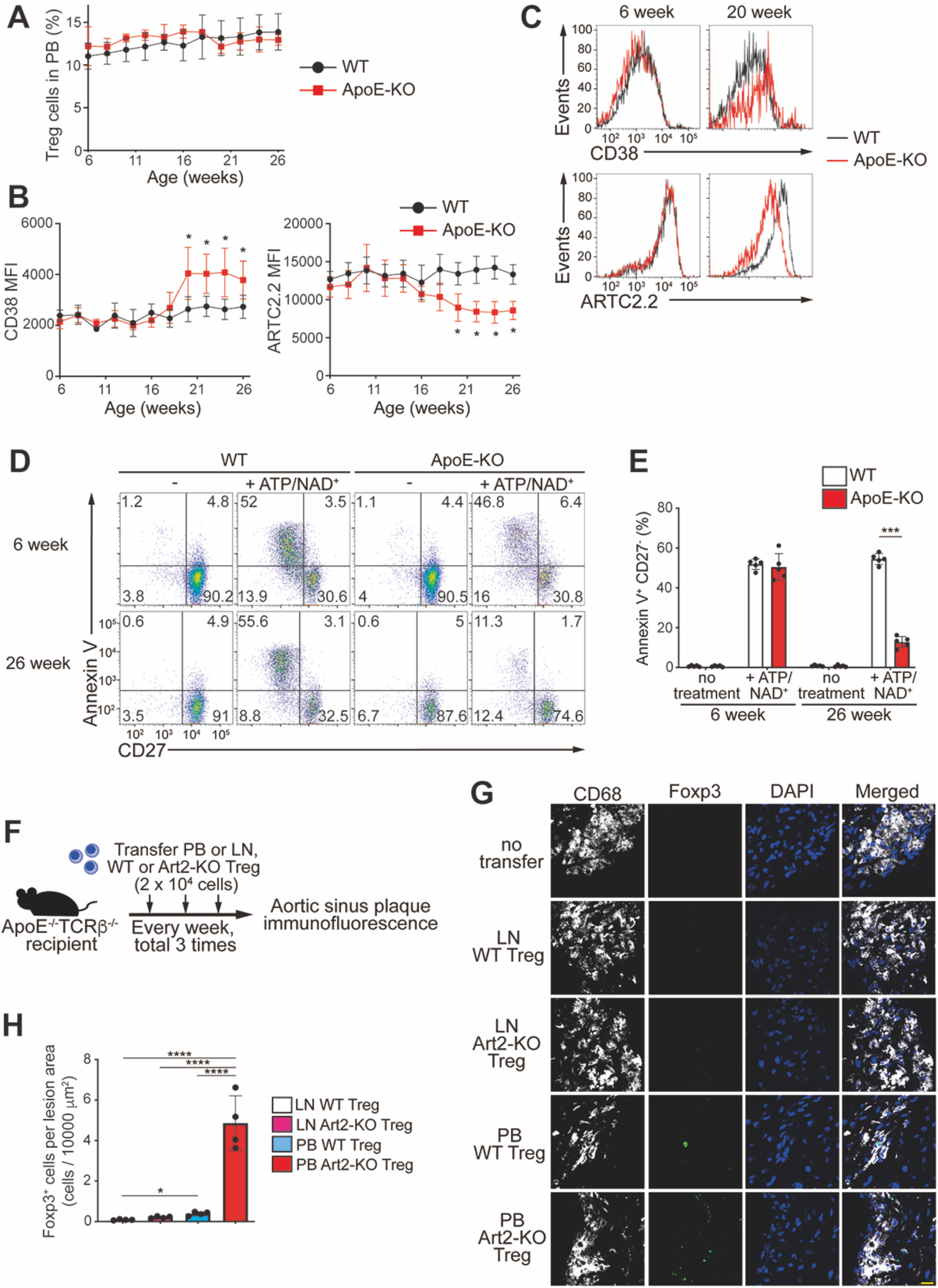
Atherosclerosis remodels PB Tregs and PB Tregs preferentially localize to plaques: **(A)** Longitudinal changes in PB Treg frequency in WT versus ApoE^−/−^ mice (n = 5 each, mean ± SD). **(B, C)** CD38 and ARTC2.2 expression levels on PB Tregs over time determined by flow cytometry (n = 5 each, mean ± SD) **(B)** and representative overlays at 6 and 20 week **(C)**. **(D, E)** Flow cytometry analysis of ATP/NAD^+^-induced apoptotic conversion of PB Tregs from WT versus ApoE^−/−^ mice at early (6 wk) and endpoint (26 wk) timepoints. Data were pooled from five independent biological replicates performed in two independent experiments, mean ± SD. **(F)** A schematic of experiments shown in **(G)**, **(H)**. **(G)** Representative plaque immunofluorescence (CD68, Foxp3, DAPI) across groups. Scale bar, 10 μm. **(H)** Quantification of **(G)**, showing Foxp3^+^ cells per lesion area. Data were pooled from four independent biological replicates performed in two independent experiments, mean ± SD. \**p* < 0.05; *** *p* < 0.005; **** *p* < 0.001, one-way ANOVA with the Bonferroni test **(B)**, **(E)**, **(H)**.

These observations further raise the hypothesis that PB Tregs are recruited to atherosclerotic plaques, where they undergo apoptosis in response to DAMP nucleotides (ATP/NAD^+^) and are subsequently engulfed by macrophages, a major cellular component of plaques. To test this model, we adoptively transferred Tregs isolated either from PB or from LN, derived from either wild-type or Art2-deficient donors, into ApoE^−/−^TCRβ^−/−^ recipients, which develop atherosclerosis despite lacking αβ T cells, and then examined donor Tregs within aortic sinus plaques (**Fig. 5F**). In recipients given wild-type donor cells, PB Tregs were detected at modestly but significantly higher numbers within plaques than LN Tregs (**Fig. 5G, 5H**). Notably, deletion of Art2—which abolishes ATP/NAD^+^ responsiveness—resulted in a pronounced increase in the number of PB-derived donor Tregs detected in plaques (**Fig. 5G, 5H**). In contrast, Art2 deficiency did not substantially increase the number of plaque-detected LN-derived Tregs (**Fig. 5G, 5H**).

Together, these data indicate that PB Tregs display enhanced tropism for atherosclerotic plaques. This preferential localization may be explained, in part, by their circulation-biased residency, which positions them to continuously patrol the vascular endothelium and access developing lesions efficiently. In addition, our tissue Treg RNA-seq analysis (**Supplementary Fig. S2**) revealed that PB Tregs express higher levels of the chemokine receptors *Ccr2* and *Cx3cr1*—receptors for CCL2 and CX3CL1, respectively, both of which are known to be upregulated within atherosclerotic lesions(Aiello *et al*, 1999; Bonacina *et al*, 2021; Combadiere *et al*, 2003)—compared with Tregs from other tissues, providing a plausible molecular basis for their increased plaque recruitment (**Supplementary Fig. S5A**).

### PB Tregs traffic to plaques and suppress plaque growth

We next assessed disease outcomes using a longer transfer regimen (weekly transfers for 14 doses starting at 9 weeks of age; **Fig. 6A**). Histologic analysis of the aortic sinus demonstrated that WT or Art2-KO PB Treg transfer significantly suppressed plaque formation compared with no-transfer controls, whereas LN Treg transfer provided weaker or no significant protection under these conditions (**Fig. 6B, 6C**). Notably, suppression of necrotic core formation was observed specifically in recipients of WT PB Tregs (**Fig. 6B, 6C**). None of the transfer conditions produced a significant change in circulating cholesterol levels (**Supplementary Fig. S5B**), indicating that the observed effects on plaque pathology were not attributable to systemic lipid lowering.

**Figure 6.**
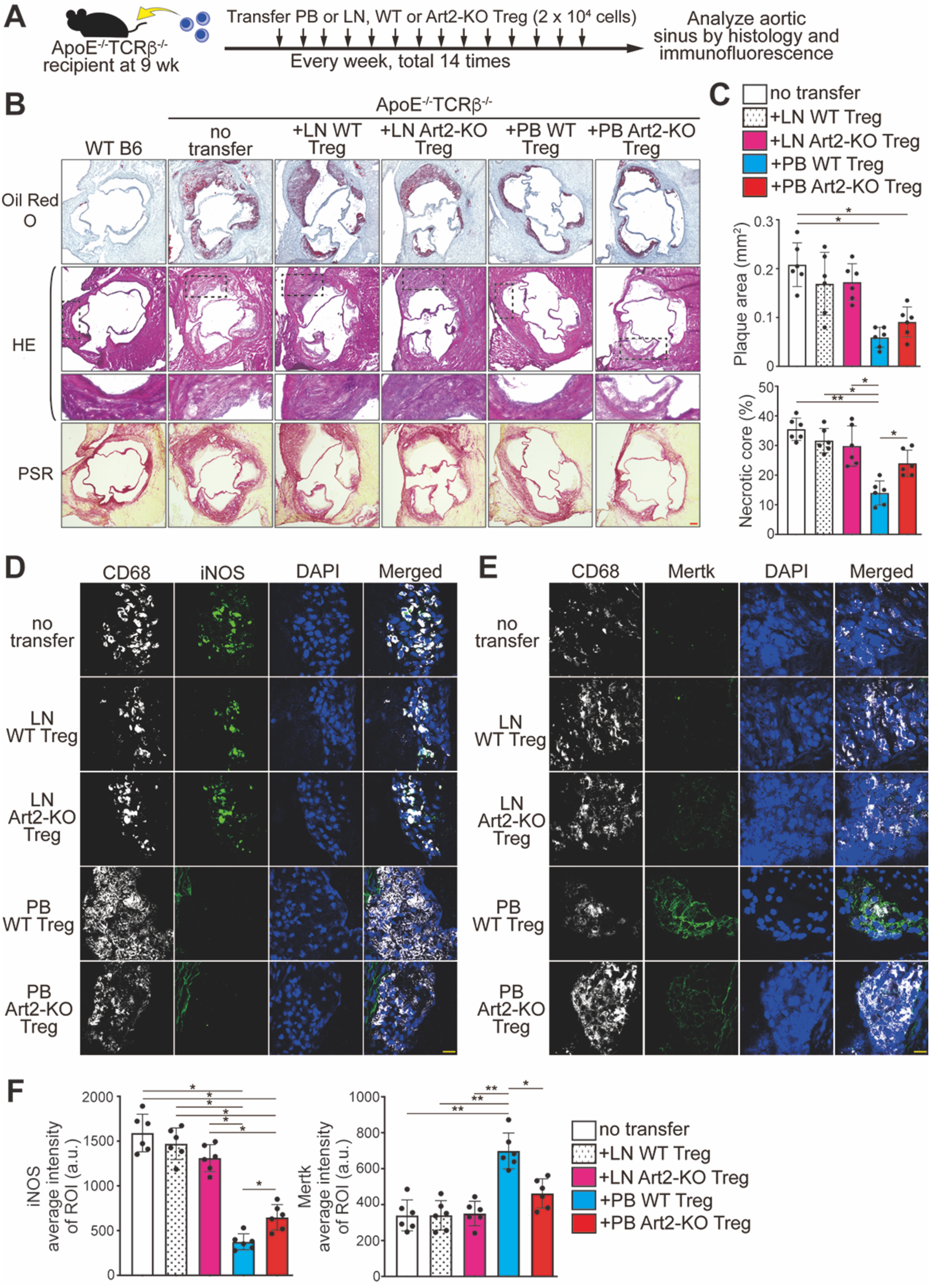
PB Tregs suppress plaque burden and necrotic core formation and modulate plaque macrophage programs: **(A)** A schematic of experiments shown in this Fig.. Long-term transfer regimen into ApoE^−/−^TCRβ^−/−^ recipients (weekly ×14 from 9 wk) followed by histology and immunofluorescence. **(B)** Representative aortic sinus serial sections, each stained with Oil Red O, Hematoxylin-Eosin (HE), and picrosirius red (PSR). For HE-stained sections, the region outlined by the dashed box is shown at higher magnification in the lower panel. Scale bar, 100 μm. **(C)** Quantification of plaque area and necrotic core as a percentage of total plaque area, based on the histologic stainings shown in **(B)**. Data were pooled from six independent biological replicates performed in three independent experiments, mean ± SD. **(D, E)** Representative immunofluorescence micrographs for CD68 with iNOS **(D)** or Mertk **(E)** and DAPI staining in aortic plaques of ApoE^-/-^TCRβ^-/-^ recipients that received the indicated cells. Scale bar, 10 μm. **(F)** Quantification of iNOS and Mertk signal intensity (arbitrary unit, a.u.) based on the immunofluorescence images in **(D)** and **(E)**, using CD68^+^ areas as regions of interest (ROI). Data were pooled from six independent biological replicates performed in three independent experiments, mean ± SD. \**p* < 0.05; ** *p* < 0.01, one-way ANOVA with the Bonferroni test **(C)**, **(F)**.

Immunofluorescence of plaques revealed that WT PB Treg transfer reduced macrophage iNOS staining and increased Mertk staining in CD68^+^ regions, consistent with suppression of inflammatory macrophage polarization and enhancement of efferocytosis-related programs (**Fig. 6D**, **6F**). Suppression of iNOS expression was also observed following transfer of Art2-KO PB Tregs; however, the magnitude of suppression was attenuated compared with that mediated by wild-type PB Tregs. (**Fig. 6D, 6F**). Together, these in vivo results indicate that PB Tregs suppress atherosclerotic plaque progression via an ARTC2.2-dependent program associated with nucleotide sensitivity and macrophage reprogramming.

### Human PB Tregs are relatively nucleotide resistant but retain P2X7R-dependent stress responses and a susceptible CD38-low subset

Finally, we asked whether analogous phenomena occur in humans. In human PBMCs, ATP/NAD^+^ treatment induced far less apoptosis than in mouse T cells, consistent with prior reports and the absence of functional ART2 in humans.(Cortes-Garcia *et al*, 2016; Haag *et al*, 1994) Nevertheless, human Tregs remained more sensitive than Tconv cells, showing increased CD27 shedding and phosphatidylserine exposure after ATP/NAD^+^ treatment; these effects were reversed by P2X7R inhibition (**Fig. 7A–7C**).

**Figure 7.**
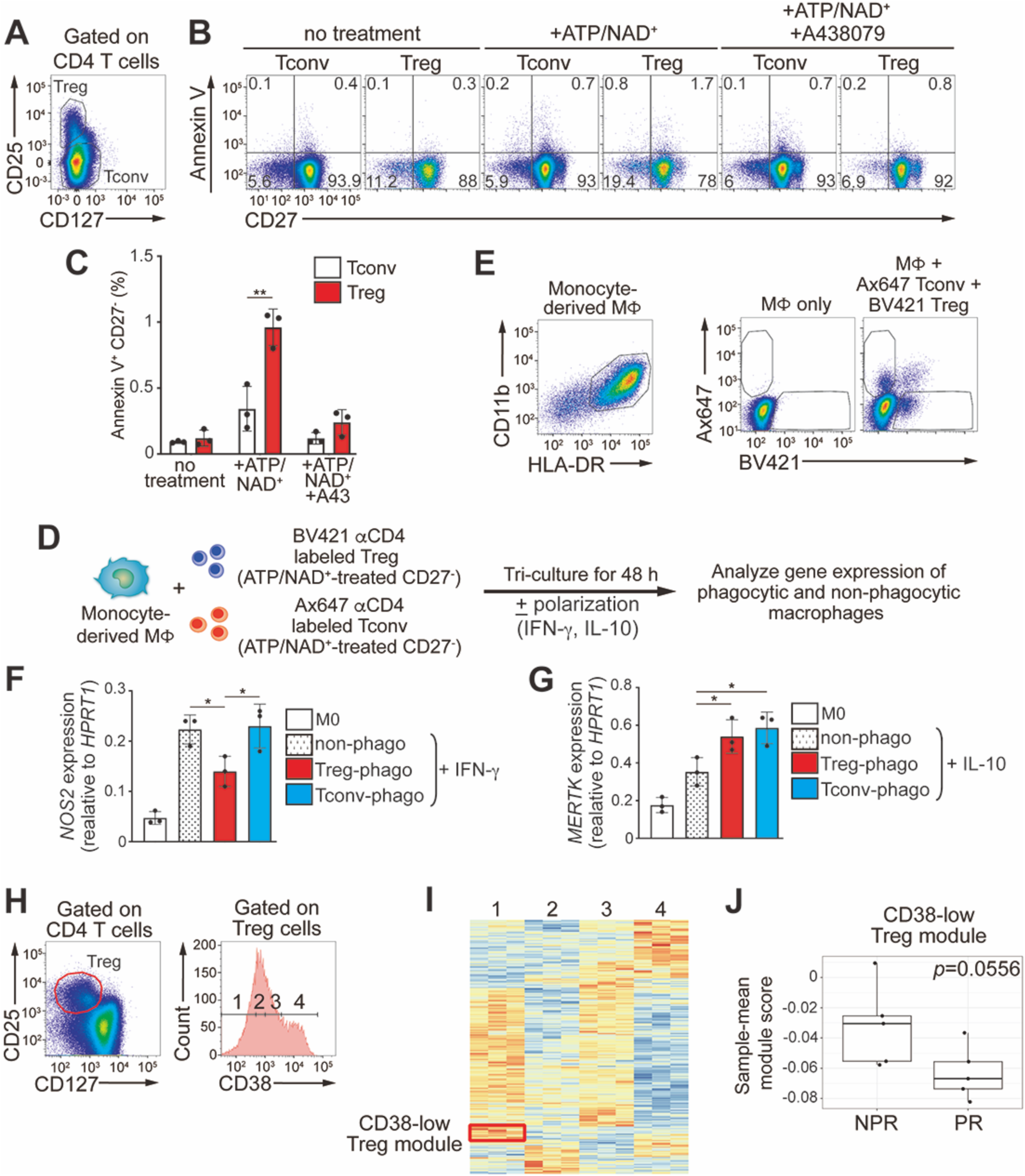
Human PB Tregs show a susceptible CD38-low fraction that is potentially reduced in plaque rupture–type acute myocardial infarction: **(A)** Gating strategy for human Tregs and Tconv cells. **(B)** Flow cytometry images of ATP/NAD^+^-induced Annexin V^+^ CD27^−^ changes in human Tregs versus Tconv, with reversal by P2X7R inhibition (A438079). **(C)** Quantification of the results shown in **(B)**. Data were pooled from three independent biological replicates, each performed with cells from different donors, mean ± SD. **(D, E)** Human monocyte-derived macrophage tri-culture scheme **(D)** and representative gating for phagocytic macrophages **(E)**. MΦ: macrophage. **(F, G)** qPCR for *NOS2* (IFN-γ polarization) **(F)** and *MERTK* (IL-10 polarization) **(G)** across macrophage subsets (non-polarized (M0), non-phagocytic, Treg-phagocytic, Tconv-phagocytic). Data are representative of three biological replicates, each performed with cells from different donors, set up in triplicate, mean ± SD. **(H)** Gating strategy for stratification of human PB Tregs into CD38 quartiles. **(I)** Heatmap of the RNA-seq result defining a gene set enriched in CD38-low Fraction 1 (“CD38-low Treg module”). **(J)** Sample-mean module score in acute myocardial infarction scRNA-seq dataset stratified by NPR (non-plaque rupture, n=5) versus PR (plaque rupture, n=5). *p*=0.556, Mann–Whitney U test. \**p* < 0.05; ** *p* < 0.01, one-way ANOVA with the Bonferroni test **(C)**, **(F)**, **(G)**.

We next tested whether efferocytosis of apoptotic human Tregs modulates macrophages. Human monocyte-derived macrophages were tri-cultured with BV421-anti-CD4-labeled Tregs and Alexa647-anti-CD4-labeled Tconv cells (both ATP/NAD^+^-treated CD27^−^ fractions), followed by IFN-γ or IL-10 polarization and sorting of phagocytic macrophages (**Fig. 7D, 7E**). In this experiment, considering the high resistance of human Treg cells, ATP/NAD^+^ treatment was performed under stronger conditions to obtain a sufficient number of cells (37°C, 30 minutes). Similar to mice, engulfment of Tregs—but not Tconv cells—suppressed IFN-γ–induced *NOS2* expression, whereas engulfment of either population enhanced IL-10–induced *MERTK* expression (**Fig. 7F, 7G**).

Because CD38 inversely correlated with nucleotide sensitivity in mouse Tregs, we stratified human PB Tregs into quartiles based on CD38 expression and challenged them with ATP/NAD^+^. The CD38-low quartile exhibited the highest nucleotide sensitivity (**Supplementary Fig. S5C–S5F**). In addition, CD38 is frequently used as an activation marker in human Tregs and has been reported to be expressed at higher levels on Tregs than on Tconv cells. Although bulk Tregs expressed higher CD38 than bulk Tconv cells (**Supplementary Fig. S5G**), the CD38-low Treg subset expressed less CD38 than the average Tconv population (**Supplementary Fig. S5H**).

Altogether, despite overall nucleotide resistance, human PB Tregs share key features with mouse PB Tregs: preferential susceptibility of a CD38-low fraction and the capacity to reprogram macrophages after efferocytosis.

### The ATP/NAD^+^-sensitive fraction of human PB Tregs is reduced in plaque rupture–type acute myocardial infarction

Collectively, our analyses in the murine atherosclerosis model revealed that progression of plaque development is accompanied by a decline in an ATP/NAD^+^-hypersensitive Treg subset, and suggested that this subset may suppress plaque formation by inducing macrophage reprogramming via efferocytosis. We therefore asked whether a comparable depletion of ATP/NAD^+^-sensitive Tregs occurs in human atherosclerotic disease. Because our experiments indicated that the CD38-low fraction of human PB Tregs exhibits increased ATP/NAD^+^ sensitivity, we stratified human PB Tregs into four quartiles based on CD38 expression (**Fig. 7H**), performed RNA-seq, and identified differentially expressed genes (DEGs) between fractions (**Supplementary Table S2**). We defined the gene set preferentially expressed in the CD38-low Fraction 1 as the “CD38-low Treg module” (**Fig. 7I**). We then leveraged this CD38-low Treg module to mine a public single-cell RNA-seq dataset from patients with acute myocardial infarction (AMI).(Qian *et al*, 2022) In this study, PB cells were profiled from patients presenting with acute myocardial infarction due to plaque rupture (PR, n=5) or non–plaque rupture (NPR; presumed plaque erosion, n=5). Because ruptured plaques are generally more macrophage-rich and necrosis-rich than eroded plaques, they are expected to exhibit heightened efferocytosis activity. Using Azimuth(Hao *et al*, 2021) to annotate and extract Tregs, we quantified the CD38-low Treg module score in PB Tregs for each sample (**Supplementary Fig. S5I-S5K**). Notably, the module score tended to be lower in PB Tregs from PR patients (*p* = 0.0556) (**Fig. 7J**), consistent with the possibility that the ATP/NAD^+^-sensitive, CD38-low PB Treg fraction is preferentially depleted in the context of plaque rupture, potentially through enhanced efferocytosis within advanced atherosclerotic lesions.

## Discussion

This study reveals that the PB Treg compartment is not simply a transient conduit between tissues. Instead, blood contains a circulation-biased Treg population enriched for low self-reactivity, low tonic TCR signaling, and a distinct epigenomic and transcriptional program. Although these PB Tregs show attenuated canonical suppressive capacity in vitro, they are uniquely poised to sense danger-associated nucleotides and to couple nucleotide-triggered apoptotic conversion to macrophage reprogramming via efferocytosis.

Tregs across tissues acquire specialized phenotypes shaped by local environmental cues, including cytokines, metabolites, and cellular interactions.(Burzyn *et al*., 2013; Feuerer *et al*., 2009; Ito *et al*., 2019) Our data suggest that weak tonic TCR signaling itself may be an important determinant of the PB Treg state. Stratification by Nur77 reporter intensity revealed a continuum in which lower tonic signaling associates with reduced CD25/IL-10 and a higher ARTC2.2/CD38 ratio, culminating in heightened ATP/NAD^+^ sensitivity. This coupling provides a conceptual framework: blood-biased, self-antigen–poor Tregs may be selected or maintained under low antigenic drive, while simultaneously becoming specialized to respond to damage-associated nucleotide cues encountered in circulation or at inflamed vascular interfaces.

A practical implication emerges from our initial observation: hypotonic RBC lysis can artifactually induce selective death of nucleotide-sensitive Treg subsets. Because ATP/NAD^+^ treatment has minimal impact on many other immune populations, this confound may have been overlooked in prior studies. Our results argue that PB immune phenotyping—especially for Tregs and other potential nucleotide-sensitive subsets—should consider RBC depletion method as an experimental variable.

Atherosclerosis engages both adaptive and innate immune responses, and prior work has established an atheroprotective role for Tregs by both depletion and adoptive transfer approaches.(Ait-Oufella *et al*., 2006; Bonacina *et al*., 2021; Klingenberg *et al*., 2013; Mor *et al*., 2007; Sharma *et al*., 2020) Mechanisms described to date largely reflect functions of viable Tregs, including modulation of macrophage polarization, promotion of efferocytosis, and, in some settings, effects on systemic lipid metabolism.(Bonacina *et al*., 2021; Proto *et al*., 2018; Sharma *et al*., 2020) Our study adds a distinct, non-canonical mechanism: PB Tregs sense plaque-associated DAMP nucleotides (ATP/NAD^+^), undergo apoptotic conversion, and reprogram macrophages upon engulfment. In ApoE^−/−^ mice, the ATP/NAD^+^-hypersensitive CD38^low^, ARTC2.2^high^ PB Treg fraction declines after plaque onset, consistent with preferential consumption of this subset; notably, the decrease begins around the reported window of aortic sinus plaque initiation and precedes accelerated plaque growth.(Reddick *et al*., 1994) Thus, our data raise the possibility that, during early atherogenesis, PB Tregs—particularly the ATP/NAD^+^-hypersensitive CD38^low^, ARTC2.2^high^ subset—protect against plaque formation through a mechanism that involves their own cell death, and that depletion of this subset may represent one trigger that accelerates disease progression. In humans, depletion of circulating Tregs has been reported in patients with vulnerable plaques,(George *et al*, 2012) and our mining of AMI scRNA-seq data suggested reduced representation of an ATP/NAD^+^-sensitive, CD38-low PB Treg program in plaque rupture—generally associated with higher macrophage content and larger necrotic cores—relative to plaque erosion. Together, these findings support a potentially conserved PB Treg–efferocytosis axis that may shape plaque inflammation and stability.

Efferocytosis is well known to promote anti-inflammatory and tissue-reparative macrophage programs, and accumulating evidence indicates that the identity of the engulfed cell can imprint distinct macrophage states.(Bosurgi *et al*, 2017; Elliott & Ravichandran, 2016; Liebold *et al*, 2024; N *et al*, 2017) Proposed mechanisms for such prey-specific imprinting include differences in the scavenger receptor pathways engaged by particular cargo and delivery of cargo-derived metabolites or other intracellular constituents that reshape macrophage signaling and transcription.(Haldar *et al*, 2014; Liebold *et al*., 2024) Using a tri-culture system that isolates phagocytic and non-phagocytic macrophages from the same microenvironment, we found that efferocytosis of apoptotic Tregs selectively dampens IFN-γ–driven inflammatory polarization while preserving IL-10–associated programs. The molecular basis for this Treg-specific suppression remains to be defined. It may involve obligate cell-surface interactions during efferocytosis—particularly because ATP/NAD^+^-treated Tregs expose phosphatidylserine yet remain viable at the time of contact (**Supplementary Fig. S4E, S4F**) —and/or delivery of Treg-enriched mediators after engulfment. For example, both mice and human Tregs contain high levels of cAMP and can transfer cAMP to other immune cells to suppress activation,(Bodor *et al*, 2012; Bopp *et al*, 2007) and cAMP signaling in macrophages can restrain inflammatory cytokine production and support alternative activation.(Luan *et al*, 2015; Wall *et al*, 2009) Consistent with the notion that Tregs harbor molecular constituents capable of directly reprogramming macrophages, Treg-derived exosomes have been reported to act directly on macrophages, suppressing M1-associated markers including iNOS while inducing M2-associated markers, with protective effects in murine models of AMI.(Hu *et al*., 2020)

Across all conditions tested—LN versus PB Tregs and wild-type versus Art2-deficient genotypes—wild-type PB Tregs conferred the most robust suppression of plaque development in adoptive transfer experiments in atherosclerosis-prone recipients lacking T cells. Enhanced efficacy of PB Tregs likely reflects their superior access to lesions: PB Tregs persist in circulation and are positioned to patrol the vascular endothelium, and they expressed higher levels of chemokine receptor genes *Ccr2* and *Cx3cr1*, which recognize CCL2 and CX3CL1, respectively—chemokines highly expressed within atherosclerotic plaques in both mice and humans.(Aiello *et al*., 1999; Bonacina *et al*., 2021; Combadiere *et al*., 2003) Consistent with this, enforced CX3CR1 expression on LN/splenic Tregs enhances plaque recruitment and atheroprotection.(Bonacina *et al*., 2021) Comparison of wild-type versus Art2-deficient PB Tregs further implicated nucleotide sensing in lesion control. Wild-type PB Tregs more strongly reduced plaque burden and, notably, necrotic core formation, whereas Art2-deficient PB Tregs provided weaker protection and did not comparably suppress necrotic core formation. These differences were mirrored at the level of plaque macrophage programs: wild-type PB Tregs reduced iNOS and increased Mertk signals within CD68^+^ regions, whereas suppression of iNOS by Art2-deficient PB Tregs was detectable but attenuated. Because iNOS contributes to tissue injury in murine and human plaques,(Buttery *et al*, 1996; Detmers *et al*, 2000; Esaki *et al*, 1997; Kockx *et al*, 2003; Kuhlencordt *et al*, 2001) and MERTK promotes efferocytosis and limits necrotic core expansion,(Cai *et al*, 2017; Qiu *et al*, 2024; Thorp *et al*, 2008) these macrophage programs provide a plausible link between the PB Treg–efferocytosis axis and plaque suppression. In addition, efferocytosis can activate LXR/PPAR transcriptional programs that induce MERTK and other phagocytic machinery, potentially creating a feed-forward loop that reinforces inflammation resolution.(Elliott *et al*, 2017; Mukundan *et al*, 2009; N *et al*, 2009)

Art2-deficient PB Tregs retained partial activity, suggesting alternative contributions from live-cell Treg mechanisms (e.g., soluble mediators represented by IL-10 and TGF-β, and contact-dependent signals).(Ouyang & Liu, 2024) Although we found that PB Tregs express relatively low levels of IL-10, they retain basal suppressive capacity, which could plausibly account for partial inhibition of iNOS induction and attenuation of plaque formation through these live-cell mechanisms. In addition, Tregs can indirectly restrain inflammatory macrophage polarization by suppressing Th1 responses. Because our in vivo experiments were performed in T cell–deficient recipients in which only Tregs were transferred, it is important to note that this experimental setting may have amplified the apparent contribution of the apoptotic PB Treg–efferocytosis axis to macrophage reprogramming relative to immune-intact atherosclerosis.

In human Tregs, overall nucleotide sensitivity was far lower than in mice Tregs as reported,(Cortes-Garcia *et al*., 2016) consistent with the lack of functional ARTC2,(Haag *et al*., 1994) yet we nonetheless detected P2X7R-dependent stress responses and identified a CD38-low Treg subset with increased susceptibility. Moreover, human macrophages were similarly reprogrammed by efferocytosis of apoptotic Tregs, supporting conservation of the downstream effector pathway. Together, these results suggest that manipulating nucleotide sensing, Treg subset composition, or efferocytosis pathways could be leveraged therapeutically—not only in atherosclerosis but potentially across inflammatory settings where extracellular ATP/NAD^+^ and macrophages are prominent.

### Study limitations

To focus on macrophage efferocytosis as a mechanistic endpoint, we performed in vivo Treg transfer experiments in ApoE^−/−^TCRβ^−/−^ recipients lacking conventional T cells. Because suppression of adaptive immune responses is an established component of atheroprotective Treg function, the relative contribution of PB Tregs in immune-intact atherosclerosis remains to be determined. In addition, IL-2 is likely limiting in T cell–deficient plaques; because IL-2 promotes Treg survival, Treg apoptosis may have been enhanced relative to immune-intact settings. Furthermore, several mechanistic questions remain open, including the molecular mediators by which engulfed Tregs selectively dampen IFN-γ polarization and the extent to which distinct PB Treg subsets are selectively recruited, converted, or depleted during disease progression. Moreover, our mining of scRNA-seq data from patients with AMI suggested a depletion of the ATP/NAD^+^-sensitive Treg subset—potentially consistent with enhanced efferocytosis in advanced plaques—but this signal did not reach statistical significance (*p* > 0.05) and was derived from a retrospective analysis with a limited sample size. Larger, prospective studies will therefore be required to validate this association in human disease.

## Data availability

ATAC-seq and RNA-seq materials availability: data reported in this paper have been deposited in the GEO database under accession number GSE325626 (tissue Treg ATAC-seq data), GSE325335 (tissue Treg RNA-seq data), GSE325329 (phagocytic BMDM RNA-seq data), and GSE325333 (human Treg RNA-seq data). Original codes generated in this study were deposited in GitHub (https://github.com/takashisekiya308/PB_Treg_Project). Any additional information needed to reanalyze the data in this paper is available from the lead contact upon request.

## Acknowledgements

We thank all the lab members or useful data discussion and acknowledge the valuable help of expert personnel at the JIHS Animal Care Facily. This work was supported by JSPS KAKENHI (Challenging Pioneering Research 23K17417), Grant for National Center for Global Health and Medicine (24A1006), and the Takeda Science Foundation.

## Author contributions

T.S. designed the research and analyzed the data; T.S., S.T., and S.H. performed the experiments; T.S. wrote the manuscript.

## Methods

### Animals

All mouse work was carried out in accordance with the guidelines for animal care approved by National Institute of Global Health and Medicine (approval # 2025-A052). Animals were maintained in specific pathogen-free conditions. Both male and female mice were used in experiments unless mentioned in the following sections. Briefly, adoptive transfer experiments were performed between sex- and age-matched mice because sex-difference are supposed to elicit immune reactions. Transcriptome and epigenomic analysis were performed with cells obtained from male mice, because sex-difference were expected to substantially affect the results of those experiments. Atherosclerosis experiments were all performed with male B6.SHL mice, because some sex differences have been detected in previous studies, with slightly more inflamed plaques shown by male mice, that conforms our interest.^(Man^ *^et al^*, ^2020)^ However, cells for flow cytometry and qRT-PCR analysis were obtained without caring sex, unless mentioned specifically, because neither differentiation pattern of cells nor expression of molecules analyzed in those studies have been reported to be affected by sex. 10-15 week old C57BL/6J Nr4a1-dsGFP mice,^(Sekiya^ *^et al.^*, ^2024)^ 12-15 week old BALB/c Nr4a1-dsGFP mice (generated by backcrossing of C57BL/6J Nr4a1-dsGFP mice to BALB/c more than five times), 10-15 week old C57BL/6J Nur77-GFP mice (016617, Jackson Laboratories),^(Moran^ *^et al.^*, ^2011)^ 10-15 week old C57BL/6J TCRβ^-/-^ (002122, Jackson Laboratories), 8-12 week old C57BL/6J Rag2^-/-^(008449, Jackson Laboratories), 10-15 week old C57BL/6J Foxp3hCD2-hCD52-KI mice (from Dr. S. Hori, RIKEN, Japan),^(Komatsu^ *^et al^*, ^2009)^ 12-15 week old BALB/c Foxp3hCD2-hCD52-KI mice (generated by backcrossing of C57BL/6J Foxp3hCD2-hCD52-KI mice to BALB/c more than five times), 10-15 week old C57BL/6J Art2a^-/-^Art2b^-/-^Foxp3hCD2-hCD52-KI mice (derived by five times backcrossing from 005347, Jackson Laboratories), and 6-10 week old B6.SHL mice^(Matsushima^ *^et al.^*, ^1999)^ were bred in NCGM’s experimental animal care facility. Breeding animals were fed ‘‘CE2’’ chow (CLEA, Japan). Experiments were performed with age-matched cohorts.

### Human samples

Approval for the human blood samples shown in this study was obtained from the Ethics Committee of the National Institute of Global Health and Medicine. Samples were from males with an age between 44 and 64 years.

### Antibodies

Monoclonal antibodies for mouse CD4 (clone L3T4, Brilliant Violet (BV) 421-, FITC-, BV510-, APC-, PE-, Alexa fluor (Ax) 488-, Ax647-, PE/Cy7-, APC/Cy7-, PE/Dazzle 594-, Pacific Blue-, or PerCP/Cy5.5-conjugated), human CD2 (clone LFA-2, Phycocerythrin (PE)/Cy7-conjugated), mouse CD45.1 (clone A20, BV510-conjugated), mouse CD45.2 (clone 104, APC/Cy7-conjugated), mouse TCR Vβ5.1/5.2 (clone MR9-4, APC-conjugated), mouse TCR Vβ11 (clone RR3-15, PE-conjugated), mouse TCR Vβ12 (clone MR11-1, PE-conjugated), mouse TCR Vβ6 (clone RR4-7, APC-conjugated), mouse TCR Vv8 (clone KJ16-133.18a, Ax647-conjugated), mouse TCR Vβ13 (clone MR12-4, PE-conjugated), mouse F4/80 (clone BM8, PE-conjugated), mouse CD38 (clone 90, APC or APC/Cy7-conjugated), mouse CD11b (clone M1/70, PE/dazzle 594-conjugated), mouse CD68 (clone FA-11, Ax647-conjugated), mouse CD25 (clone PC61, BV510-conjugated), human CD25 (clone BC96, PE-conjugated), human CD4 (clone RPA-T4, PerCP/Cy5.5-conjugated), human CD127 (clone A019D5, APC/Cy7-conjugated), human HLA-DR (clone Tu39, APC/Fire 750-conjugated), and human CD38 (clone HB-7, BV510-conjugated) were purchased from BioLegend. Monoclonal antibodies for mouse Foxp3 (clone FJK-16s, Ax488-conjugated), mouse/human CD27 (clone LG.7F9, APC-conjugated), mouse iNOS (clone CXNFT, Ax488-conjugated), and mouse Mertk (clone DS5MMER, Ax488-conjugated) were purchased from Thermo Fisher Scientific. Monoclonal antibodies for human CD11b (clone ICRF44, PE-conjugated) was purchased from BD Pharmingen. Monoclonal antibodies for mouse ARTC2.2 (clone Nika102, PE-conjugated) was purchased from Novus Biologicals. Unconjugated anti-mouse CD3ε (clone 2C11) and anti-mouse CD28 (clone 57.31) antibodies were purchased from BioLegend.

### Red blood cell removal from mouse peripheral blood and flow cytometry

Red blood cells (RBCs) were removed from mouse peripheral blood by either hypotonic lysis or density-gradient centrifugation. For RBC lysis, whole blood was mixed with five volumes of RBC lysis buffer (0.15 M NH_4_Cl, 10 mM KHCO_3_, 0.2 mM EDTA), gently inverted for 1 min, centrifuged at 4,000 rpm for 1 min, and the supernatant was discarded. The cell pellet was used as RBC-depleted peripheral blood cells. For density-gradient separation, 1 mL of blood was diluted with 4 mL PBS, layered onto 4 mL Lymphoprep (Veritas, ST-18060), and centrifuged at 800 × g for 35 min at 20°C; cells at the interphase were collected as RBC-depleted peripheral blood cells. Cells were washed to remove residual reagents and stained for flow cytometry. For apoptosis assessment, cells were stained for 15 min at room temperature in Annexin V binding buffer (10 mM HEPES, pH 7.4; 140 mM NaCl; 2.5 mM CaCl_2_) with PE-conjugated Annexin V (BioLegend, 640907) and antibodies against CD4, human CD2 (Foxp3 reporter), and CD27, then analyzed by flow cytometry. In selected experiments, whole blood was pre-incubated with the P2X7 receptor inhibitor A438079 (Selleck, S7705; 40 µM) or an anti-ARTC2.2 neutralizing nanobody (BioLegend, 149802; 20 µg/mL) throughout the whole steps.

### Autologous mixed lymphocyte reaction (MLR)

Lymph nodes- and peripheral blood-derived Treg cells (CD4^+^Foxp3^+^) were isolated from Nr4a1-dsGFP;Foxp3-hCD2-hCD52-KI double-reporter mice and used as responder cells. lymph nodes Tregs from Foxp3-hCD2-hCD52 knock-in single-reporter mice were used as non-transgenic controls for the Nr4a1-dsGFP reporter. Responder cells (2 × 10^4^) were rested for 48 h in RPMI 1640 supplemented with 10% FBS, IL-2 (40 ng/mL; PeproTech 212-12), and 55 µM β-mercaptoethanol to reduce baseline GFP. Responder cells were then co-cultured for 24 h either with T cell–depleted syngeneic splenocytes (2 × 10^4^) or under plate-bound anti-CD3 (1 µg/mL) plus soluble anti-CD28 (0.5 µg/mL) stimulation. Nr4a1-dsGFP fluorescence in responder cells was assessed by flow cytometry.

### ATAC-seq

ATAC-seq was performed using Nextera DNA Library Preparation Kit (illumina). 4×10^5^ cells (male origin) were suspended in 200 μl Hypotonic buffer (20 mM Tris pH 7.5, 10 mM NaCl, 3 mM), and incubated for 15 min on ice. Then, 10 μl 10% NP-40 was added to the sample and vortexed for 10 sec at maximum speed. Cells were pelleted by centrifugation at 6,000 rpm for 5 min, then suspended in 40 μl 1x Tagmentation buffer. Samples were added with 2 μl Tagment DNA Enzyme 1, and incubated for 30 min at 37 °C. Then, samples were cleaned up with FastGene Gel/PCR clean up kit (Nippon Genetics), and eluted with 20 μl elution buffer. 5 μl of the eluted samples were PCR amplified with index 1 (i7) and index 2 (i5) adapter primers, using PCR Primer Cocktail which was attached to Nextera DNA Library Preparation Kit. 150 bp paired end next generation sequencing was performed with HiSeq X Ten (illumina). Generated sequencing reads in FASTQ format were mapped to mm10 reference genome using bowtie2. Reads mapped to the “black list” regions were removed with intersectBed command, inputting the bed file which denotes the black list region (mm10-blacklist.v2.bed.gz).

### Adoptive transfer into Rag2^−/−^ mice to assess circulation-biased Tregs

Sorted lymph nodes CD45.2^+^ total CD4^+^ T cells (1 × 10^6^) derived from Foxp3 reporter mice (CD45.2^+^ Foxp3-hCD2-hCD52-KI) were mixed with sorted CD45.1^+^ Tregs (1 × 10^4^) derived from either lymph nodes or peripheral blood of Foxp3 reporter mice (CD45.1^+^ Foxp3-hCD2-hCD52-KI) and intravenously transferred into 8–12-week-old Rag2^− / −^ recipients. One week after transfer, cells were isolated from recipient peripheral blood and lymph nodes, and donor-derived Tregs were identified by CD4, CD45.1, and human CD2 (Foxp3 reporter) staining.

### ATP/NAD^+^ treatment of mouse PB Tregs

RBC-depleted peripheral blood cells obtained by density-gradient separation were incubated with ATP (10 µM; Sigma-Aldrich A7699) plus NAD^+^ (3 µM; Sigma-Aldrich N11636) for 10 min at room temperature. Where indicated, a CD38 inhibitor (Sigma-Aldrich 5.38763; 50 nM) was included. After washing, cells were stained in Annexin V binding buffer with PE-Annexin V and antibodies against CD4, human CD2 (Foxp3 reporter), and CD27 for 15 min at room temperature, followed by flow cytometry. For macrophage phagocytosis experiments, Annexin V was avoided because it can inhibit efferocytosis. Instead, after ATP/NAD^+^ treatment, cells were stained in FACS buffer (PBS containing 2 mM EDTA and 0.5% BSA) with antibodies against CD4, hCD2, and CD27, and CD27^-^ cells were purified by sorting for subsequent assays.

### Preparation of single-cell suspensions from mouse tissues

Single-cell suspensions from lymph nodes, thymus, and spleen were prepared by mechanical dissociation through a 70-µm cell strainer (Corning). Bone marrow cells were flushed from tibiae and femora using a 26G needle and syringe, followed by RBC lysis. For intestine, liver, and lung, tissues were processed (e.g., removal of intestinal contents), minced, and digested in RPMI 1640 + 10% FBS containing DNase I (1 mg/mL) and collagenase D (0.5 mg/mL) for 45 min at 37°C. Digests were filtered through a 100-µm strainer, centrifuged (2,000 rpm, 5 min), and pellets were subjected to a Percoll gradient (37% over 70%) followed by centrifugation (2,000 rpm, 25 min, room temperature). Cells at the interphase were collected.

### In vitro suppression assay

Responder cells (Ly5.1^+^CD4^+^CD25^low^Foxp3^-^; 2×10^4^ cells) from Ly5.1^+^ C57/BL6 Foxp3-hCD2-hCD52-KI reporter mice were labeled with 5 μM CellTrace Far Red (Invitrogen, C34564), then cultured with or without sorted Treg cells (CD4^+^ Foxp3^+^) from Ly5.2^+^ C57/BL6 Foxp3-hCD2-hCD52-KI mice at the indicated ratio for 96 hr at 37°C in 200 μl of RPMI1640 medium plus 10% FBS, supplemented with 1 μg/ml anti-CD3 antibodies (clone 2C11) and 55 μM β-mercaptoethanol, in the presence of 2×10^4^ T cell-depleted splenocytes irradiated at 20 Gy, in round-bottomed 96-well culture plates. Responder proliferation was quantified by dilution of CellTrace Far Red in Ly5.1^+^ cells by flow cytometry.

### Calcein-AM–based live/dead assessment

ATP/NAD^+^-treated CD45.1^+^ PB cells and untreated CD45.2^+^ PB cells (both from Foxp3 reporter mice) were mixed 1:1 and stained with Annexin V, CD4, CD45.2, and hCD2 in Annexin V binding buffer. Cells were then resuspended in 100 nM Calcein-AM in Annexin V binding buffer and immediately analyzed by flow cytometry. As a dead-cell control, LN cells were heat-treated at 70°C for 30 min.

### RNA extraction and qRT-PCR

Total RNA was extracted using RNAiso PLUS (Takara, 9109). cDNA was synthesized with ReverTra Ace qPCR RT Master Mix with gDNA Remover (Toyobo, FQS-301). qPCR was performed on a StepOne Real-Time PCR System (Applied Biosystems) using THUNDERBIRD SYBR qPCR Mix (Toyobo, QPS-101). Primer sequences are available upon request.

### Bulk RNA-seq and differential expression analysis

Total RNA was extracted with RNAiso PLUS (TAKARA, 9109). Samples were further cleaned using an NucleoSpin RNA Clean-up XS (MACHEREY-NAGEL). Then, RNA-seq libraries were prepared with NEBNext Ultra II Directional RNA Library Prep kit for Illumina (NEB, E7760), using Multiplex Oligos for Illumina (Dual Index Primers Set 1, NEB, E7600), following the manufacturer’s protocol. Next generation sequencing were performed using Illumina Hiseq X or Illumina NovaSeq6000 with a standard 150-bp paired-end read protocol. Fastq output files were aligned to mm10 reference genome with STAR(Dobin *et al*, 2013), then quantification of gene expression were calculated with RSEM.(Li & Dewey, 2011) Differentially expressed genes were identified by DESeq2.(Love *et al*, 2014) Principal component analysis was performed with the “plotPCA” function on the data processed with the “variance stabilizing transformation (vsd)” method using DESeq2. Euclidian distances between samples were performed on vst-normalized matrix using “dist(method=“euclidean”) function of R.

### Bone marrow–derived macrophages (BMDMs) and phagocytosis assays (mouse)

Bone marrow cells from wild-type C57BL/6 mice were isolated from tibiae and femora, subjected to RBC lysis, and cultured at 1 × 10^6^/mL in high-glucose DMEM + 10% FBS containing M-CSF (10 ng/mL) for 7 days; fresh medium was added on day 4. On day 7, BMDMs (1 x 10^6^) were co-cultured with ATP/NAD^+^-treated CD27^−^ Tconv or Treg cells (2.5 x 10^5^ each) labeled with fluorophore-conjugated anti-CD4 antibodies (Alexa Fluor 647 or BV421) for 48 h, in the presence or absence of polarizing cytokines IFN-γ (BioLegend #575304, 50 ng/mL, 48 h) or IL-10 (BioLegend #575804, 50 ng/mL, 48 h). Macrophages were harvested and sorted as CD11b^+^F4/80^+^ cells, with phagocytic status inferred from Alexa647/BV421 positivity. For Annexin V blockade experiments, ATP/NAD^+^-treated CD27^−^ Treg cells were pre-incubated with recombinant Annexin V (BD Pharmingen, 556416; 10 μg/mL) for 10 min at room temperature prior to mixing with BMDMs. Annexin V (10 μg/mL) was maintained in the culture medium throughout the subsequent 3-h co-culture period.

### Longitudinal flow cytometric profiling of PB Tregs in WT and ApoE-deficient mice

Peripheral blood was serially collected from the same cohort of wild-type (WT) and ApoE-deficient (ApoE-KO) mice (both on the Foxp3–hCD2–hCD52-KI background) every other week from 6 to 26 weeks of age. Red blood cells were removed by hypotonic lysis, and leukocytes were stained with antibodies against CD4, hCD2, ARTC2.2, and CD38 for flow cytometric analysis. PB Tregs were identified as CD4^+^hCD2^+^ cells, and surface expression of CD38 and ARTC2.2 was quantified as mean fluorescence intensity (MFI) at each time point.

### Treg transfer into atherosclerosis model mice, aortic sinus histology, and plasma cholesterol

ApoE^−/−^TCRβ^−/−^ mice received weekly intravenous injections of 2 × 10^4^ Tregs (CD4^+^Foxp3 reporter^+^) derived from PB or LNs of wild-type or Art2-deficient donors (both Foxp3-hCD2-hCD52-KI background), starting at 9 weeks of age and continuing for 14 weeks. Recipient mice were maintained on normal chow (CLEA, CE2). One week after the final transfer, mice were perfused with 15 mL PBS and hearts were dissected and embedded in OCT (Sakura Finetek) and frozen. Serial 10-µm sections were prepared across the ∼400-µm aortic sinus region and distributed across five slides for Oil Red O, H&E, Picrosirius Red, iNOS immunostaining, and MERTK immunostaining. Picrosirius Red staining was performed using a Picro-Sirius Red Stain Kit for Collagen (ScyTek Laboratories, PSR-1). Plaque area and necrotic core size were quantified using ImageJ. Necrotic core regions were defined primarily on H&E by absence of nuclei and eosinophilic debris (often with clefts), corroborated by Picrosirius Red (fibrous cap with strong signal overlying a relatively weakly stained core) and supported by Oil Red O (adjacent foam-cell–rich regions). Total plasma cholesterol was measured from 1 µL plasma using LabAssay Cholesterol (Wako, 293-93601).

### Immunofluorescence staining of frozen sections

Frozen tissue sections prepared as described above were subjected to immunofluorescence staining for Foxp3 and iNOS. Sections were fixed in pre-chilled 100% methanol (−20°C) for 10 min, briefly air-dried, and rehydrated in Tris-buffered saline (TBS) (3 × 10 min). Sections were circumscribed with a hydrophobic barrier pen and blocked for 30 min at room temperature in blocking buffer containing 10% normal serum, 1% BSA, and 0.05% Tween-20. Fluorophore-conjugated primary antibodies against Foxp3 and iNOS (diluted in TBS containing 1% BSA, 0.1% cold fish skin gelatin, 0.5% Triton X-100, and 0.05% sodium azide) were incubated overnight at 4°C. Sections were washed in TBS containing 0.05% Tween-20 (3 × 10 min), counterstained with DAPI for 1 min, washed again in TBS (3 × 10 min), and mounted using an antifade mounting medium. For Mertk staining, sections were fixed in pre-chilled acetone (−20°C) for 5 min, briefly air-dried, and rehydrated in PBS (2 × 3 min). After circumscribing and blocking as above, sections were incubated with a fluorophore-conjugated anti-MERTK antibody overnight at 4°C. Sections were washed in TBS containing 0.05% Tween-20 (3 × 10 min with gentle agitation), counterstained with DAPI for 1 min, washed in TBS (3 × 10 min), and mounted with antifade mounting medium. Images were acquired using an Olympus FV-1000 confocal microscope. For quantitative comparisons, imaging settings (laser power, detector gain/offset, pinhole, and scan speed) were kept constant across experimental groups within each experiment, and representative fields were acquired using identical acquisition parameters. Fluorescence quantification was performed in Fluoview (Olympus) software using predefined regions of interest (ROIs). When macrophage-restricted quantification was required, CD68^+^ areas were used to define ROIs, within which iNOS or MERTK signal intensity was measured after background subtraction. Image acquisition and/or downstream quantification were performed in a blinded manner to experimental condition whenever feasible, and multiple sections per animal and multiple fields per section were analyzed as indicated in the corresponding Fig. legends.

### Human peripheral blood Treg and Tconv cell isolation and ATP/NAD^+^ stimulation

Human Tregs (CD4^+^CD25^high^CD127^low^) Tconv cells (non-Treg CD4^+^ T cells) were sorted from PBMCs obtained from buffy coats prepared from 10–20 mL peripheral blood of healthy donors by density-gradient centrifugation with Lymphoprep (Veritas, ST-18060). The study was approved by the Institutional Review Board of the National Institute for Global Health and Medicine and conducted in accordance with the Declaration of Helsinki; written informed consent was obtained from all donors. Sorted Tregs were stimulated with ATP (10 µM) plus NAD^+^ (3 µM) for 10 min at room temperature, washed, stained with PE-Annexin V and antibodies against CD4 and CD27 in Annexin V binding buffer for 15 min at room temperature, and analyzed by flow cytometry. For phagocytosis experiments requiring higher cell yields, ATP/NAD^+^ stimulation was performed at 37°C for 30 min as indicated.

### Human monocyte-derived macrophages and phagocytosis assays

PBMCs were isolated from 10–20 mL peripheral blood from three donors by Lymphoprep. CD14+ monocytes were enriched by positive selection using CD14 MicroBeads (Miltenyi, 130-087-052). Cells were cultured at 1 × 10^6^/mL in RPMI 1640 supplemented with 10% human AB serum (Sigma-Aldrich, H6914) and either GM-CSF (50 ng/mL) for subsequent IFN-γ polarization or M-CSF (50 ng/mL) for subsequent IL-10 polarization for 6 days, with half-medium exchange on day 3. On day 6, autologous Tregs and Tconv cells were isolated from the same donor, labeled with BV421- or Alexa647-conjugated anti-CD4 antibodies, treated with ATP/NAD^+^, and CD27− cells were purified by sorting. Macrophages were co-cultured with these T cells and polarized with IFN-γ (50 ng/mL, 48 h) or IL-10 (50 ng/mL, 48 h). Macrophages were then sorted as CD11b^+^HLA-DR^+^ cells, with phagocytic status determined by BV421/Alexa647 positivity.

### Assessment of external scRNA-seq data sets

For single-cell analysis of the human myocardial infarction peripheral blood mononuclear cells, the “barcodes.tsv.gz”, “features.tsv.gz”, and “matrix.mtx.gz” files of each patient’s sample was downloaded from GSE269269.(Qian *et al*., 2022) Downstream analyses were performed using the Seurat R package(Hao *et al*., 2021) to regress out the mitochondrial contents and cell cycle, to SCT-normalize the scores, and reduce dimensionality. Cell lineages were determined with Azimuth R package(Hao *et al*., 2021), employing the “pbmcref” as a reference. To derive the expression scores of the “CD38-low Treg module”, the gene list (n=26, shown in **Supplementary Table 2**) was incorporated into the Seurat subject with an “AddModuleScore” function. Expression scores of the the “CD38-low Treg module” were analyzed for the “Treg” subset.

### Quantification and statistical analysis

p values were calculated with Graphpad Prism software (for ANOVA tests and Student’s t test) or R (for Mann–Whitney U test). p values of less than 0.05 were considered significant. All error bars in graphs represent SEM calculated at least three replicates. Data were assessed for normal distribution and plotted in the Fig.s as mean ±SD. No samples or animals were excluded from the analyses.

## References

Adriouch S, Hubert S, Pechberty S, Koch-Nolte F, Haag F, Seman M (2007) NAD+ released during inflammation participates in T cell homeostasis by inducing ART2-mediated death of naive T cells in vivo. J Immunol 179: 186–194

Aiello RJ, Bourassa PA, Lindsey S, Weng W, Natoli E, Rollins BJ, Milos PM (1999) Monocyte chemoattractant protein-1 accelerates atherosclerosis in apolipoprotein E-deficient mice. Arterioscler Thromb Vasc Biol 19: 1518–1525

Ait-Oufella H, Salomon BL, Potteaux S, Robertson AK, Gourdy P, Zoll J, Merval R, Esposito B, Cohen JL, Fisson S et al (2006) Natural regulatory T cells control the development of atherosclerosis in mice. Nat Med 12: 178–180

Aswad F, Kawamura H, Dennert G (2005) High sensitivity of CD4+CD25+ regulatory T cells to extracellular metabolites nicotinamide adenine dinucleotide and ATP: a role for P2X7 receptors. J Immunol 175: 3075–3083

Bodor J, Bopp T, Vaeth M, Klein M, Serfling E, Hunig T, Becker C, Schild H, Schmitt E (2012) Cyclic AMP underpins suppression by regulatory T cells. Eur J Immunol 42: 1375–1384

Bonacina F, Martini E, Svecla M, Nour J, Cremonesi M, Beretta G, Moregola A, Pellegatta F, Zampoleri V, Catapano AL et al (2021) Adoptive transfer of CX3CR1 transduced-T regulatory cells improves homing to the atherosclerotic plaques and dampens atherosclerosis progression. Cardiovasc Res 117: 2069–2082

Bopp T, Becker C, Klein M, Klein-Hessling S, Palmetshofer A, Serfling E, Heib V, Becker M, Kubach J, Schmitt S et al (2007) Cyclic adenosine monophosphate is a key component of regulatory T cell-mediated suppression. J Exp Med 204: 1303–1310

Bosurgi L, Cao YG, Cabeza-Cabrerizo M, Tucci A, Hughes LD, Kong Y, Weinstein JS, Licona-Limon P, Schmid ET, Pelorosso F et al (2017) Macrophage function in tissue repair and remodeling requires IL-4 or IL-13 with apoptotic cells. Science 356: 1072–1076

Burzyn D, Kuswanto W, Kolodin D, Shadrach JL, Cerletti M, Jang Y, Sefik E, Tan TG, Wagers AJ, Benoist C et al (2013) A special population of regulatory T cells potentiates muscle repair. Cell 155: 1282–1295

Buttery LD, Springall DR, Chester AH, Evans TJ, Standfield EN, Parums DV, Yacoub MH, Polak JM (1996) Inducible nitric oxide synthase is present within human atherosclerotic lesions and promotes the formation and activity of peroxynitrite. Lab Invest 75: 77–85

Cai B, Thorp EB, Doran AC, Sansbury BE, Daemen MJ, Dorweiler B, Spite M, Fredman G, Tabas I (2017) MerTK receptor cleavage promotes plaque necrosis and defective resolution in atherosclerosis. J Clin Invest 127: 564–568

Chinetti-Gbaguidi G, Colin S, Staels B (2015) Macrophage subsets in atherosclerosis. Nat Rev Cardiol 12: 10–17

Combadiere C, Potteaux S, Gao JL, Esposito B, Casanova S, Lee EJ, Debre P, Tedgui A, Murphy PM, Mallat Z (2003) Decreased atherosclerotic lesion formation in CX3CR1/apolipoprotein E double knockout mice. Circulation 107: 1009–1016

Cortes-Garcia JD, Lopez-Lopez C, Cortez-Espinosa N, Garcia-Hernandez MH, Guzman-Flores JM, Layseca-Espinosa E, Portales-Cervantes L, Portales-Perez DP (2016) Evaluation of the expression and function of the P2X7 receptor and ART1 in human regulatory T-cell subsets. Immunobiology 221: 84–93

Detmers PA, Hernandez M, Mudgett J, Hassing H, Burton C, Mundt S, Chun S, Fletcher D, Card DJ, Lisnock J et al (2000) Deficiency in inducible nitric oxide synthase results in reduced atherosclerosis in apolipoprotein E-deficient mice. J Immunol 165: 3430–3435

Dobin A, Davis CA, Schlesinger F, Drenkow J, Zaleski C, Jha S, Batut P, Chaisson M, Gingeras TR (2013) STAR: ultrafast universal RNA-seq aligner. Bioinformatics 29: 15–21

Elliott MR, Koster KM, Murphy PS (2017) Efferocytosis Signaling in the Regulation of Macrophage Inflammatory Responses. J Immunol 198: 1387–1394

Elliott MR, Ravichandran KS (2016) The Dynamics of Apoptotic Cell Clearance. Dev Cell 38: 147–160

Esaki T, Hayashi T, Muto E, Yamada K, Kuzuya M, Iguchi A (1997) Expression of inducible nitric oxide synthase in T lymphocytes and macrophages of cholesterol-fed rabbits. Atherosclerosis 128: 39–46

Feuerer M, Herrero L, Cipolletta D, Naaz A, Wong J, Nayer A, Lee J, Goldfine AB, Benoist C, Shoelson S et al (2009) Lean, but not obese, fat is enriched for a unique population of regulatory T cells that affect metabolic parameters. Nat Med 15: 930–939

George J, Schwartzenberg S, Medvedovsky D, Jonas M, Charach G, Afek A, Shamiss A (2012) Regulatory T cells and IL-10 levels are reduced in patients with vulnerable coronary plaques. Atherosclerosis 222: 519–523

Haag F, Koch-Nolte F, Kuhl M, Lorenzen S, Thiele HG (1994) Premature stop codons inactivate the RT6 genes of the human and chimpanzee species. J Mol Biol 243: 537–546

Haldar M, Kohyama M, So AY, Kc W, Wu X, Briseno CG, Satpathy AT, Kretzer NM, Arase H, Rajasekaran NS et al (2014) Heme-mediated SPI-C induction promotes monocyte differentiation into iron-recycling macrophages. Cell 156: 1223–1234

Hansson GK, Hermansson A (2011) The immune system in atherosclerosis. Nat Immunol 12: 204–212

Hao Y, Hao S, Andersen-Nissen E, Mauck WM, 3rd, Zheng S, Butler A, Lee MJ, Wilk AJ, Darby C, Zager M et al (2021) Integrated analysis of multimodal single-cell data. Cell 184: 3573–3587 e3529

Herbin O, Ait-Oufella H, Yu W, Fredrikson GN, Aubier B, Perez N, Barateau V, Nilsson J, Tedgui A, Mallat Z (2012) Regulatory T-cell response to apolipoprotein B100-derived peptides reduces the development and progression of atherosclerosis in mice. Arterioscler Thromb Vasc Biol 32: 605–612

Hsieh CS, Zheng Y, Liang Y, Fontenot JD, Rudensky AY (2006) An intersection between the self-reactive regulatory and nonregulatory T cell receptor repertoires. Nat Immunol 7: 401–410

Hu H, Wu J, Cao C, Ma L (2020) Exosomes derived from regulatory T cells ameliorate acute myocardial infarction by promoting macrophage M2 polarization. IUBMB Life 72: 2409–2419

Hubert S, Rissiek B, Klages K, Huehn J, Sparwasser T, Haag F, Koch-Nolte F, Boyer O, Seman M, Adriouch S (2010) Extracellular NAD+ shapes the Foxp3+ regulatory T cell compartment through the ART2-P2X7 pathway. J Exp Med 207: 2561–2568

Ito M, Komai K, Mise-Omata S, Iizuka-Koga M, Noguchi Y, Kondo T, Sakai R, Matsuo K, Nakayama T, Yoshie O et al (2019) Brain regulatory T cells suppress astrogliosis and potentiate neurological recovery. Nature 565: 246–250

Jordan MS, Boesteanu A, Reed AJ, Petrone AL, Holenbeck AE, Lerman MA, Naji A, Caton AJ (2001) Thymic selection of CD4+CD25+ regulatory T cells induced by an agonist self-peptide. Nat Immunol 2: 301–306

Josefowicz SZ, Lu LF, Rudensky AY (2012) Regulatory T cells: mechanisms of differentiation and function. Annu Rev Immunol 30: 531–564

Klingenberg R, Gerdes N, Badeau RM, Gistera A, Strodthoff D, Ketelhuth DF, Lundberg AM, Rudling M, Nilsson SK, Olivecrona G et al (2013) Depletion of FOXP3+ regulatory T cells promotes hypercholesterolemia and atherosclerosis. J Clin Invest 123: 1323–1334

Kockx MM, Cromheeke KM, Knaapen MW, Bosmans JM, De Meyer GR, Herman AG, Bult H (2003) Phagocytosis and macrophage activation associated with hemorrhagic microvessels in human atherosclerosis. Arterioscler Thromb Vasc Biol 23: 440–446

Komatsu N, Mariotti-Ferrandiz ME, Wang Y, Malissen B, Waldmann H, Hori S (2009) Heterogeneity of natural Foxp3+ T cells: a committed regulatory T-cell lineage and an uncommitted minor population retaining plasticity. Proc Natl Acad Sci U S A 106: 1903–1908

Kuhlencordt PJ, Chen J, Han F, Astern J, Huang PL (2001) Genetic deficiency of inducible nitric oxide synthase reduces atherosclerosis and lowers plasma lipid peroxides in apolipoprotein E-knockout mice. Circulation 103: 3099–3104

Levine AG, Arvey A, Jin W, Rudensky AY (2014) Continuous requirement for the TCR in regulatory T cell function. Nat Immunol 15: 1070–1078

Li B, Dewey CN (2011) RSEM: accurate transcript quantification from RNA-Seq data with or without a reference genome. BMC Bioinformatics 12: 323

Liebold I, Al Jawazneh A, Casar C, Lanzloth C, Leyk S, Hamley M, Wong MN, Kylies D, Grafe SK, Edenhofer I et al (2024) Apoptotic cell identity induces distinct functional responses to IL-4 in efferocytic macrophages. Science 384: eabo7027

Love MI, Huber W, Anders S (2014) Moderated estimation of fold change and dispersion for RNA-seq data with DESeq2. Genome Biol 15: 550

Luan B, Yoon YS, Le Lay J, Kaestner KH, Hedrick S, Montminy M (2015) CREB pathway links PGE2 signaling with macrophage polarization. Proc Natl Acad Sci U S A 112: 15642–15647

Maganto-Garcia E, Tarrio ML, Grabie N, Bu DX, Lichtman AH (2011) Dynamic changes in regulatory T cells are linked to levels of diet-induced hypercholesterolemia. Circulation 124: 185–195

Mallat Z, Ait-Oufella H, Tedgui A (2007) Regulatory T-cell immunity in atherosclerosis. Trends Cardiovasc Med 17: 113–118

Man JJ, Beckman JA, Jaffe IZ (2020) Sex as a Biological Variable in Atherosclerosis. Circ Res 126: 1297–1319

Matsushima Y, Hayashi S, Tachibana M (1999) Spontaneously hyperlipidemic (SHL) mice: Japanese wild mice with apolipoprotein E deficiency. Mamm Genome 10: 352–357

Moore KJ, Sheedy FJ, Fisher EA (2013) Macrophages in atherosclerosis: a dynamic balance. Nat Rev Immunol 13: 709–721

Mor A, Planer D, Luboshits G, Afek A, Metzger S, Chajek-Shaul T, Keren G, George J (2007) Role of naturally occurring CD4+ CD25+ regulatory T cells in experimental atherosclerosis. Arterioscler Thromb Vasc Biol 27: 893–900

Moran AE, Holzapfel KL, Xing Y, Cunningham NR, Maltzman JS, Punt J, Hogquist KA (2011) T cell receptor signal strength in Treg and iNKT cell development demonstrated by a novel fluorescent reporter mouse. J Exp Med 208: 1279–1289

Mukundan L, Odegaard JI, Morel CR, Heredia JE, Mwangi JW, Ricardo-Gonzalez RR, Goh YP, Eagle AR, Dunn SE, Awakuni JU et al (2009) PPAR-delta senses and orchestrates clearance of apoptotic cells to promote tolerance. Nat Med 15: 1266–1272

N AG, Bensinger SJ, Hong C, Beceiro S, Bradley MN, Zelcer N, Deniz J, Ramirez C, Diaz M, Gallardo G et al (2009) Apoptotic cells promote their own clearance and immune tolerance through activation of the nuclear receptor LXR. Immunity 31: 245–258

N AG, Quintana JA, Garcia-Silva S, Mazariegos M, Gonzalez de la Aleja A, Nicolas-Avila JA, Walter W, Adrover JM, Crainiciuc G, Kuchroo VK et al (2017) Phagocytosis imprints heterogeneity in tissue-resident macrophages. J Exp Med 214: 1281–1296

Ouyang X, Liu Z (2024) Regulatory T cells and macrophages in atherosclerosis: from mechanisms to clinical significance. Front Immunol 15: 1435021

Proto JD, Doran AC, Gusarova G, Yurdagul A, Jr., Sozen E, Subramanian M, Islam MN, Rymond CC, Du J, Hook J et al (2018) Regulatory T Cells Promote Macrophage Efferocytosis during Inflammation Resolution. Immunity 49: 666–677 e666

Qian J, Gao Y, Lai Y, Ye Z, Yao Y, Ding K, Tong J, Lin H, Zhu G, Yu Y et al (2022) Single-Cell RNA Sequencing of Peripheral Blood Mononuclear Cells From Acute Myocardial Infarction. Front Immunol 13: 908815

Qiu S, Liu J, Chen J, Li Y, Bu T, Li Z, Zhang L, Sun W, Zhou T, Hu W et al (2024) Targeted delivery of MerTK protein via cell membrane engineered nanoparticle enhances efferocytosis and attenuates atherosclerosis in diabetic ApoE(-/-) Mice. J Nanobiotechnology 22: 178

Reddick RL, Zhang SH, Maeda N (1994) Atherosclerosis in mice lacking apo E. Evaluation of lesional development and progression. Arterioscler Thromb 14: 141–147

Rivas-Yanez E, Barrera-Avalos C, Parra-Tello B, Briceno P, Rosemblatt MV, Saavedra-Almarza J, Rosemblatt M, Acuna-Castillo C, Bono MR, Sauma D (2020) P2X7 Receptor at the Crossroads of T Cell Fate. Int J Mol Sci 21

Sakaguchi S, Mikami N, Wing JB, Tanaka A, Ichiyama K, Ohkura N (2020) Regulatory T Cells and Human Disease. Annu Rev Immunol 38: 541–566

Sekiya T, Hidano S, Takaki S (2024) Tonic TCR and IL-1beta signaling mediate phenotypic alterations of naive CD4(+) T cells. Cell Rep 43: 113954

Sekiya T, Kashiwagi I, Yoshida R, Fukaya T, Morita R, Kimura A, Ichinose H, Metzger D, Chambon P, Yoshimura A (2013) Nr4a receptors are essential for thymic regulatory T cell development and immune homeostasis. Nat Immunol 14: 230–237

Seman M, Adriouch S, Scheuplein F, Krebs C, Freese D, Glowacki G, Deterre P, Haag F, Koch-Nolte F (2003) NAD-induced T cell death: ADP-ribosylation of cell surface proteins by ART2 activates the cytolytic P2X7 purinoceptor. Immunity 19: 571–582

Sharma M, Schlegel MP, Afonso MS, Brown EJ, Rahman K, Weinstock A, Sansbury BE, Corr EM, van Solingen C, Koelwyn GJ et al (2020) Regulatory T Cells License Macrophage Pro-Resolving Functions During Atherosclerosis Regression. Circ Res 127: 335–353

Tanaka S, Maeda S, Hashimoto M, Fujimori C, Ito Y, Teradaira S, Hirota K, Yoshitomi H, Katakai T, Shimizu A et al (2010) Graded attenuation of TCR signaling elicits distinct autoimmune diseases by altering thymic T cell selection and regulatory T cell function. J Immunol 185: 2295–2305

Tang Q, Bluestone JA (2013) Regulatory T-cell therapy in transplantation: moving to the clinic. Cold Spring Harb Perspect Med 3

Thorp E, Cui D, Schrijvers DM, Kuriakose G, Tabas I (2008) Mertk receptor mutation reduces efferocytosis efficiency and promotes apoptotic cell accumulation and plaque necrosis in atherosclerotic lesions of apoe-/- mice. Arterioscler Thromb Vasc Biol 28: 1421–1428

Wall EA, Zavzavadjian JR, Chang MS, Randhawa B, Zhu X, Hsueh RC, Liu J, Driver A, Bao XR, Sternweis PC et al (2009) Suppression of LPS-induced TNF-alpha production in macrophages by cAMP is mediated by PKA-AKAP95-p105. Sci Signal 2: ra28

Wigren M, Bjorkbacka H, Andersson L, Ljungcrantz I, Fredrikson GN, Persson M, Bryngelsson C, Hedblad B, Nilsson J (2012) Low levels of circulating CD4+FoxP3+ T cells are associated with an increased risk for development of myocardial infarction but not for stroke. Arterioscler Thromb Vasc Biol 32: 2000–2004

Witztum JL, Lichtman AH (2014) The influence of innate and adaptive immune responses on atherosclerosis. Annu Rev Pathol 9: 73–102

Yazdani M, Khosropanah S, Hosseini A, Doroudchi M (2016) Resting and Activated Natural Tregs Decrease in the Peripheral Blood of Patients with Atherosclerosis. Iran J Immunol 13: 249–262

