## Supplementary figures for "Low-self-reactive, circulation-biased blood regulatory T cells sense danger-associated nucleotides to restrain atherosclerosis"

### Supplementary Information

Supplementary Information contains Supplementary Figures S1 to S5.

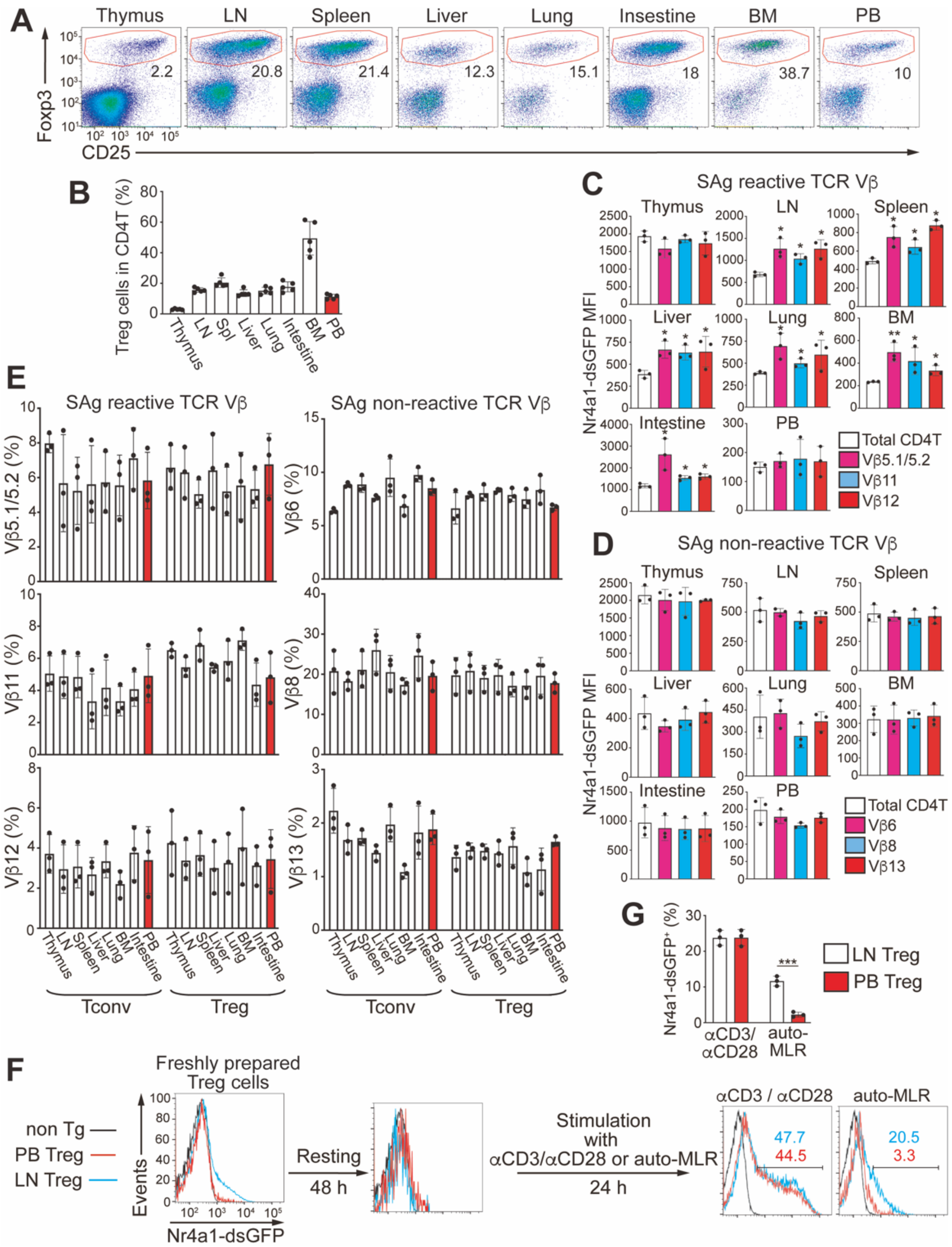


**Supplementary Figure S1. Peripheral blood Tregs are enriched for cells with reduced self-reactivity.** **(A)** Flow cytometric profiles of Foxp3 and CD25 expression in CD4 T cells across tissues. Numbers adjacent to outlined areas indicate percent Treg cells. **(B)** Quantification of the results in **(A)**. Treg frequency among CD4 T cells across tissues are shown. Each dot represents an individual mouse, mean ± SD (n = 5). LN; lymph nodes, Spl: spleen, BM: bone marrow; PB: peripheral blood. **(C, D)** Nr4a1-dsGFP mean fluorescence intensities (MFI) in Tregs with superantigen-reactive **(C)** and non-reactive **(D)** TCR Vβ subsets across tissues of BALB/c mice. Each dot represents an individual mouse, mean ± SD (n = 3). **(E)** Frequencies of superantigen-reactive and non-reactive Vβ subsets in Tconv versus Tregs across tissues (C57BL6). Each dot represents an individual mouse, mean ± SD (n = 3). **(F)** Flow cytometric profiles show autologous MLR workflow and Nr4a1-dsGFP induction in PB versus LN Tregs after αCD3/αCD28 or auto-MLR stimulation. Numbers adjacent to outlined areas indicate percent $GFP^+$ cells. **(G)** Quantification of the results in **(F)**. **p* < 0.05; ****p*<0.005, one-way ANOVA with the Bonferroni test **(C)** and Student's t test **(G)**.

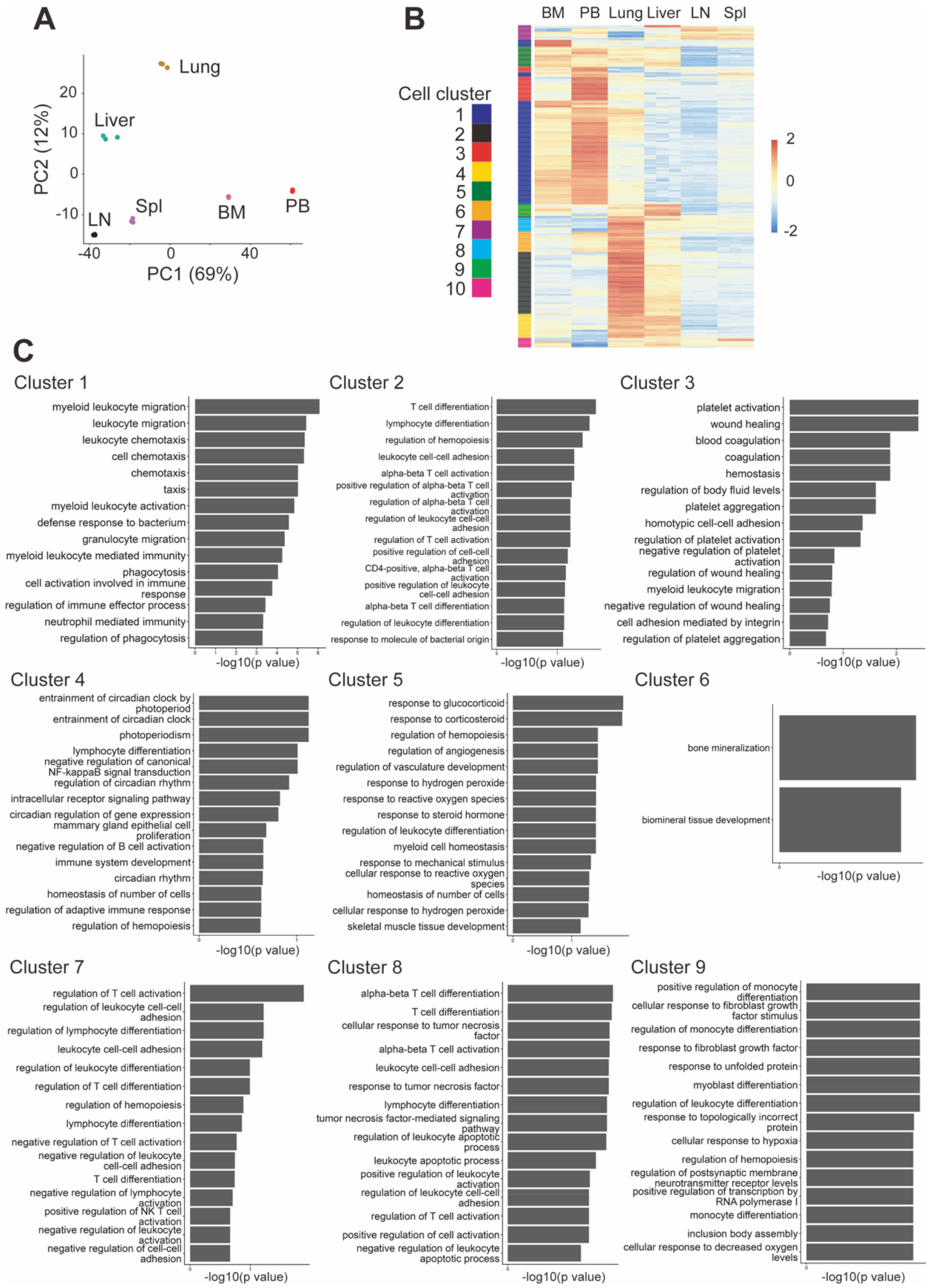


**Supplementary Figure S2. PB Tregs exhibit a distinct transcriptional program.**
**(A)** Principal component analysis of bulk RNA-seq profiles of tissue Tregs isolated from the indicated organs. BM: bone marrow; LN: lymph nodes; PB: peripheral blood; Spl: spleen. **(B)** Heatmap of the RNA-seq result showing tissue-specific gene clusters across tissues. **(C)** Gene ontology (GO) enrichment for representative clusters identified in **(B)**.

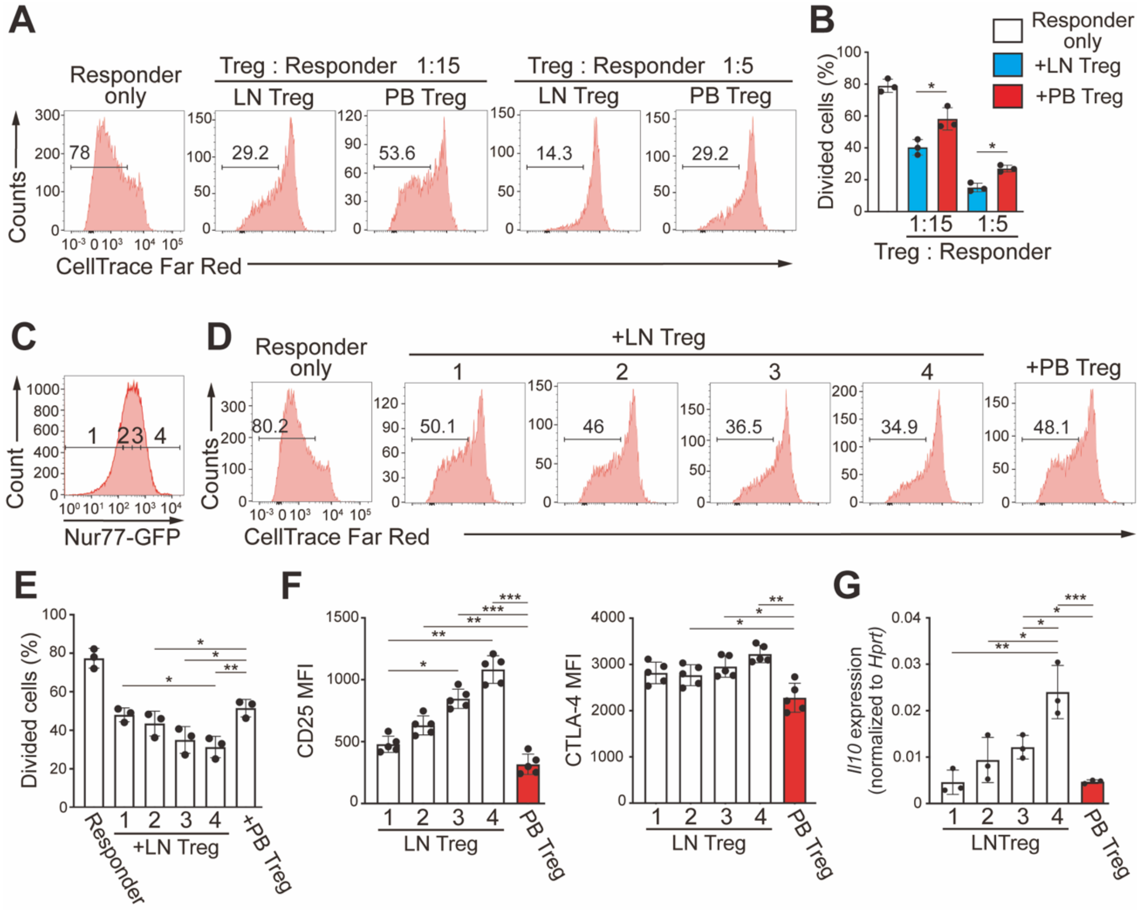


**Supplementary Figure S3. PB Tregs show reduced suppressive capacity, linked to low tonic TCR signaling and reduced suppressor-associated factors.**

**(A)** In vitro suppression assay of conventional CD4 T cells (Responder) labeled with CellTrace Far Red, in the presence or absence of lymph nodes (LN) or peripheral blood (PB) Tregs at indicated ratios. Numbers indicate the percentage of cells in the indicated area. **(B)** Quantification of the results in **(A)**. **(C)** Nur77-GFP intensity–based stratification of LN Tregs into four fractions (1–4). **(D)** In vitro suppression assay with each LN Treg fraction (fractions 1–4, depicted in **(C)**) and comparison to PB Tregs. **(E)** Quantification of the results in **(D)**. **(F, G)** CD25 and CTLA-4 mean fluorescence intensity (MFI) **(F)**, and *Il10* mRNA expression **(G)** across LN Treg fractions (fractions 1–4, depicted in **(C)**) and PB Tregs. Each dot in **(F)** represents an individual mouse, mean ± SD (n = 5). Results shown in **(B)**, **(E)** and **(G)** are representative of three biological replicates performed in three independent experiments, set up in triplicate, mean ± SD. *$p < 0.05$; **$p < 0.01$; ***$p < 0.005$, one-way ANOVA with the Bonferroni test **(B)**, **(E)**, **(F)** and **(G)**.

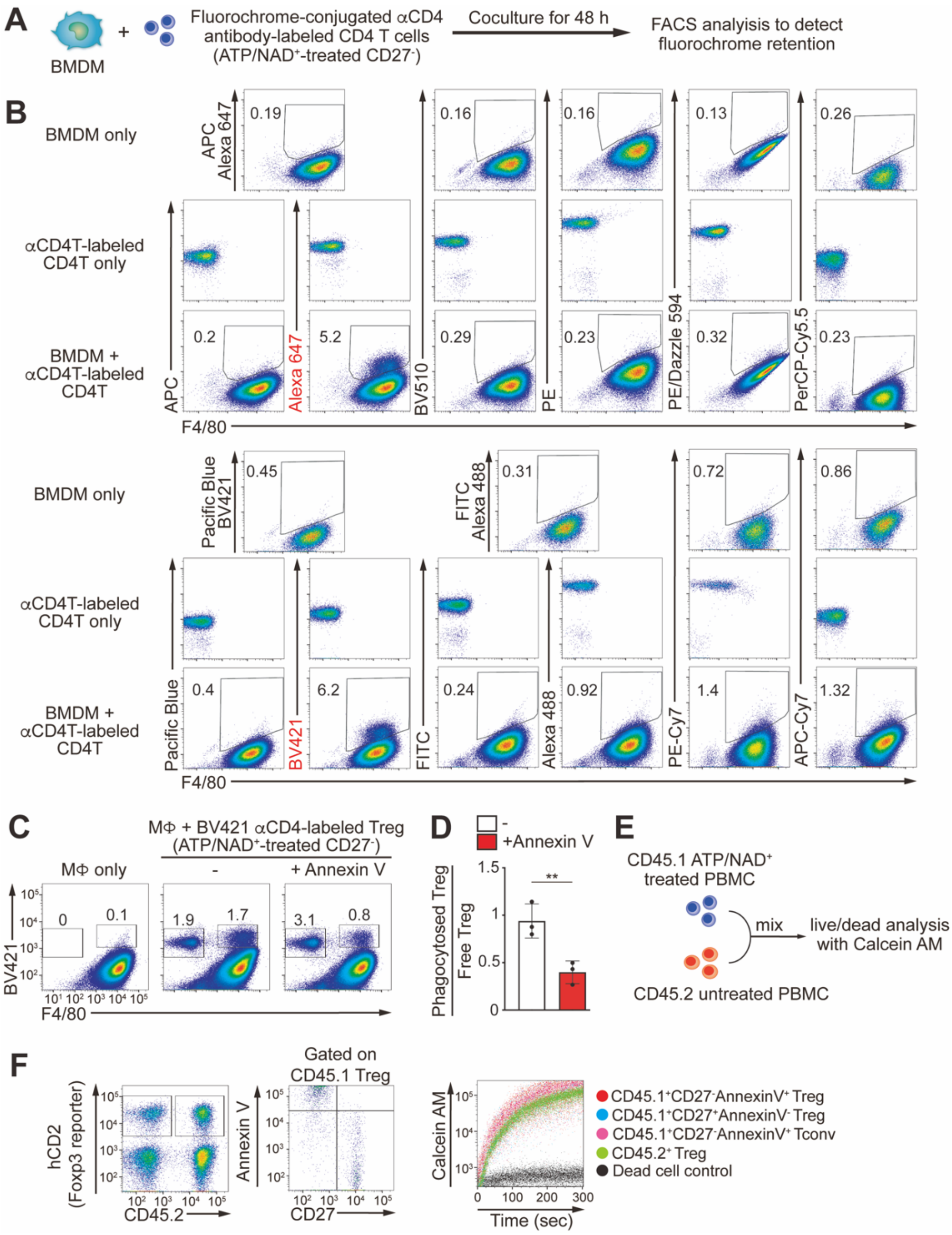


**Supplementary Figure S4. Characterization of apoptotic Treg cell phagocytosis by bone marrow-derived macrophages.**
**(A)** Screening scheme for fluorochrome retention in bone marrow-derived macrophages (BMDM) after uptake of anti-CD4-labeled T cells. **(B)** Representative flow cytometry plots showing durable retention of phagocytosed CD4 T cell-derived Brilliant Violet 421 and Alexa fluor 647 signals in BMDMs after 48 h of coculture. **(C, D)** Annexin V blockade reduces uptake of ATP/NAD$^+$-treated CD27$^-$ Tregs; representative plots **(C)** and quantification **(D)**. ATP/NAD$^+$-treated, CD27$^-$ Treg cells were mixed with BMDMs and analyzed by flow cytometry after 3 hours of incubation. Data are representative of three biological replicates, set up in triplicate, mean ± SD. **(E)** Strategy for the live/dead determination of ATP/NAD$^+$-treated CD4 T cells performed in **(F)**. **(F)** Left and Middle: Gating strategy to identify ATP/NAD$^+$-untreated (CD45.1$^-$) and -treated (CD45.1$^+$) Tconv and Treg cells which were either undergoing (Annexin V$^+$ CD27$^-$) or not undergoing (Annexin V$^-$ CD27$^+$) apoptotic conversion. Right: A representative flow plot of Calcein-AM analysis, in which live cells incorporate and hydrolyze Calcein-AM thus become fluorescent. $^{**}p < 0.01$, Student's t test **(D)**.

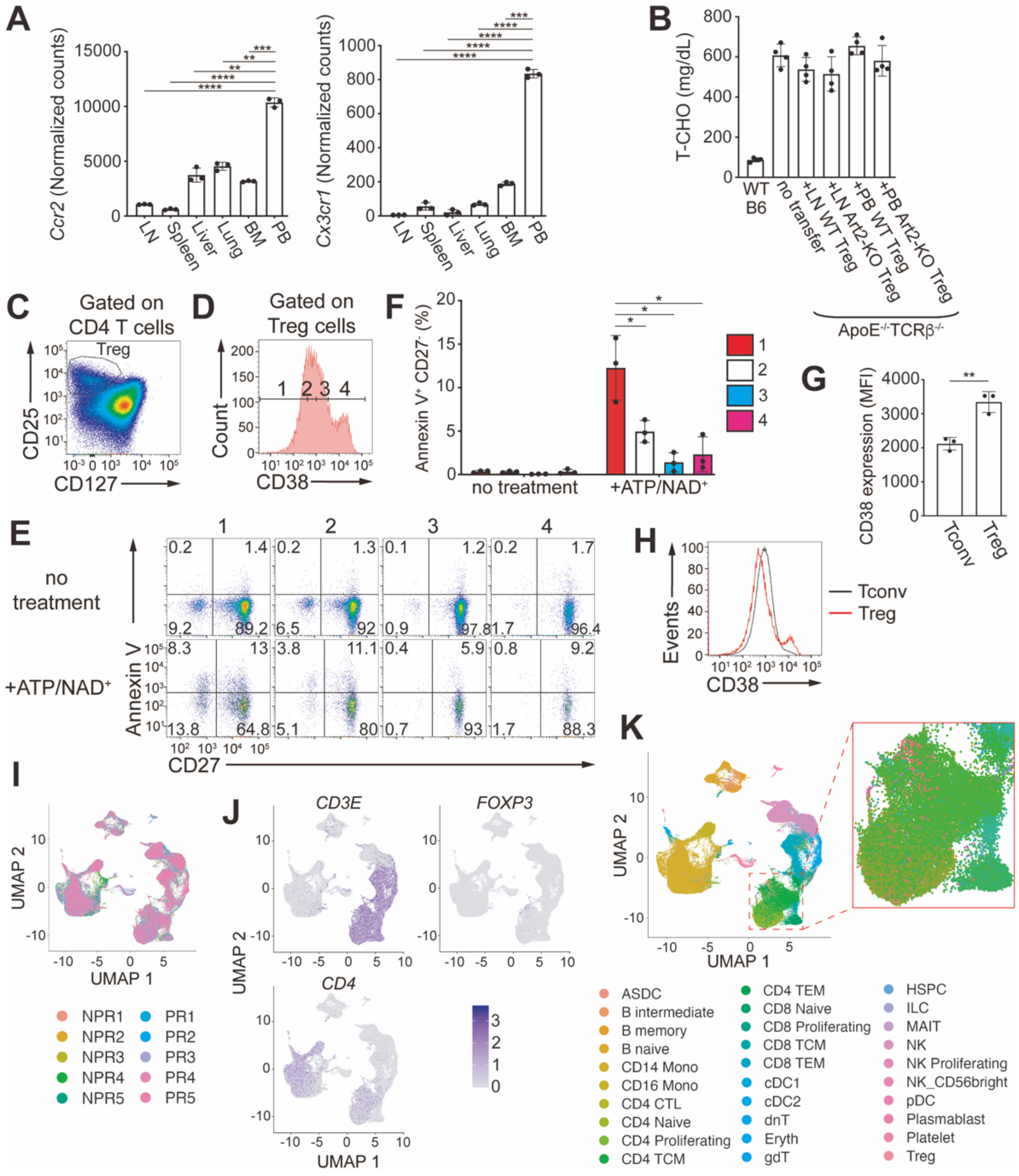


**Supplementary Figure S5. Plaque tropism and atheroprotective roles of peripheral blood Tregs.** **(A)** RNA-seq results of *Ccr2* and *Cx3cr1* expression across tissue Tregs. **(B)** Total plasma cholesterol (T-CHO) levels across wild-type (WT) and ApoE$^{-/-}$TCRβ$^{-/-}$recipients with indicated transfers. Each dot represents an individual mouse, mean ± SD (n = 4). **(C, D)** Gating strategy for human Treg isolation **(C)** and further stratification into CD38 quartiles **(D)**. **(E)** Flow cytometry shows representative Annexin V versus CD27 plots for each fraction defined in **(D)** under the indicated ATP/NAD$^+$ stimulation conditions. **(F)** Quantification of cells undergoing apoptotic conversion (Annexin V$^+$ CD27$^-$) in **(E)**. Data are representative of three biological replicates, each performed with three independent donors, set up in triplicate, mean ± SD. **(G)** Bulk CD38 expression in human PB Tregs versus Tconv cells. CD38 mean fluorescence intensity (MFI) was compared between bulk Tregs and bulk conventional CD4 T cells. Each dot represents an individual donor, mean ± SD (n = 3). **(H)** Representative flow cytometry overlays of CD38 expression on bulk Tregs and Tconv cells. **(I–K)** UMAP visualizations of the acute myocardial infarction scRNA-seq dataset used for module-score analysis in **Figure 7**, showing distribution of cells from independent donors **(I)**, marker genes expression **(J)**, and cell subset identities **(K)**. For Treg plot, the region outlined by the dashed box is shown at higher magnification in the right panel. *$p < 0.05$; ** $p < 0.01$; *** $p < 0.005$; **** $p < 0.001$, one-way ANOVA with the Bonferroni test **(A)**, **(G)**, **(F)**.
