## Supplementary Table S1 for "Low-self-reactive, circulation-biased blood regulatory T cells sense danger-associated nucleotides to restrain atherosclerosis"

| Supplementary Table S1, Genes assigned to the expression clusters in tissue-derived Treg cells | | | | | | | | | |
|---|---|---|---|---|---|---|---|---|---|
| Cluster 1 | Cluster 2 | Cluster 3 | Cluster 4 | Cluster 5 | Cluster 6 | Cluster 7 | Cluster 8 | Cluster 9 | Cluster 10 |
| 6430548M08Rik | Aebp2 | Adgre5 | Areg | Adam8 | Acer2 | Actb | Abcb1a | Atf4 | Cish |
| Abca9 | Aff1 | Alox15 | Atp2b4 | Adrb2 | Acod1 | Dtx1 | Adamts6 | Btg1 | Cxcr3 |
| Ace | Aff4 | Ampd3 | Bhlhe40 | Anxa1 | Antxr2 | Ecm1 | Bzw2 | Cd69 | Gpr68 |
| Adgre1 | Arhgap31 | Apoe | Ccr4 | Anxa2 | Atp11b | Gpr83 | Cbarp | Dnajb1 | Maf |
| Adgre4 | Arid5a | Asah1 | Cenpa | Aqp1 | Atp2b1 | Gramd1a | Cd2 | Hsp90aa1 | Myb |
| Adipor1 | Bcl2l11 | Bach1 | Cpm | Atf3 | Cd53 | Hspa8 | Cd83 | Itgb1 | Nav2 |
| Adpgk | Birc3 | Cald1 | Crip1 | Btg2 | Dennd4a | Hsph1 | Dusp4 | Jun | Penk |
| Adss1 | Bltp1 | Cbl | Csrnp1 | Ccr2 | Emb | Ift80 | Icos | Junb | Ptma |
| Aldh2 | C9orf72 | Cd9 | Ctla4 | Cd44 | Ern1 | Il2ra | Itk | Lars2 | Rpl10a |
| Anxa3 | Cblb | Ctla2a | Dusp5 | Chil3 | Ets2 | Lag3 | Lrig1 | Nr4a1 | Rplp1 |
| Apobec1 | Ccr7 | Ctss | Gadd45b | Cited2 | Fosl2 | Lat | Nfkb1 | Ppp1r15a | Rps3 |
| Apobr | Chd7 | Ednrb | Gata3 | Cxcl2 | Gpcpd1 | Limd2 | Nr4a3 | Rgs16 | Rrm2 |
| App | Cnbp | Entpd1 | Gem | Cxcr4 | Il18rap | Ltb | Nt5e | S100a10 | Serpina3g |
| Arap3 | Crbn | Gm2a | Gimap6 | Dstn | Irs2 | Macf1 | Pdcd1 | S100a4 | Tbc1d4 |
| Asprv1 | Crlf3 | Gp5 | Glrx | Dusp1 | Lpl | Msl1 | Plagl1 | Slc38a2 | Ttn |
| Atp1a3 | Crybg1 | Gzma | H3f3b | Fgl2 | Map3k1 | Neb | Rundc3b | Tamalin | |
| Bst1 | Cytip | Il17ra | Hexim1 | Fos | Mmp19 | Nsg2 | Stat5a | Ubc | |
| C3 | D16Ertd472e | Il6ra | Hmgb2 | Fosb | Mxd1 | Pfn1 | Tnfrsf18 | Zfp36l1 | |
| C5ar1 | Dgat1 | Itgb2 | Id2 | Gadd45a | Mxi1 | Ppp1r9b | Tnfrsf4 | | |
| Card9 | Dusp10 | Itgb5 | Ifngr1 | H1f2 | Pgm2l1 | Ptpn6 | Tnfrsf9 | | |
| Ccl6 | Eea1 | Klf3 | Ifrd1 | Hbb-b1 | Rap1b | Ptpn7 | Tnfsf8 | | |
| Ccl9 | Elf1 | Lcp1 | Il18r1 | Klf2 | Rgs2 | Rasal3 | Traf1 | | |
| Ccpg1 | Ell2 | Mapk14 | Il1rl1 | Klf4 | Sbno2 | Tcf7 | | | |
| Cd14 | Elovl5 | Mcl1 | Il7r | Klf6 | Sgk1 | | | | |
| Cd177 | Epas1 | Mef2a | Irf2bp2 | Rhob | Srgn | | | | |
| Cd19 | Epb41 | Mfap3l | Lamc1 | S100a11 | Ssh2 | | | | |
| Cd244a | Ets1 | Mmrn1 | Nfkbia | Serinc3 | Tcp11l2 | | | | |
| Cd24a | Ezr | Mpl | Nr4a2 | Trib1 | Tent5a | | | | |
| Cd300a | Fam107b | Ms4a4c | Per1 | Tsc22d3 | Tpm4 | | | | |
| Cd300ld | Fus | Mylk | Pim1 | Vim | Trim25 | | | | |
| Cd300lf | Gabpb1 | Nav1 | Rgs1 | Zfp36 | Tubb2a | | | | |
| Cd36 | Gna13 | Notch2 | Rora | Zfp36l2 | | | | | |
| Cd68 | Got1 | P2ry12 | Sik1 | | | | | | |
| Cd74 | Gpr132 | Prg4 | Sla | | | | | | |
| Cd79b | Gramd2b | Ptpn12 | Spty2d1 | | | | | | |
| Cfp | H2az1 | Pttg1ip | Tgif1 | | | | | | |
| Ckap4 | Hif1a | Rnf149 | Tnfaip3 | | | | | | |
| Clec12a | Hivep2 | Sell | | | | | | | |
| Clec4a1 | Ifnar1 | Sh3bgrl2 | | | | | | | |
| Clec4d | Ikzf3 | Slc2a3 | | | | | | | |
| Clec4e | Il4ra | Slc6a4 | | | | | | | |
| Clec5a | Isy1 | Treml1 | | | | | | | |
| Col1a2 | Itgav | Tsc22d1 | | | | | | | |
| Csf1r | Itpkb | Txnip | | | | | | | |
| Csf3r | Jak2 | Zfp710 | | | | | | | |
| Cst3 | Jmy | Zyx | | | | | | | |
| Ctsd | Kdm2b | | | | | | | | |
| Ctsh | Kdm6b | | | | | | | | |
| Cx3cr1 | Lemd3 | | | | | | | | |
| Cybb | Madd | | | | | | | | |
| Cyp4f18 | Nabp1 | | | | | | | | |
| Dennd3 | Neurl3 | | | | | | | | |
| Dhrs9 | Nfil3 | | | | | | | | |
| Dmxl2 | Nfkb2 | | | | | | | | |
| Dock5 | Nfkbiz | | | | | | | | |
| Dusp6 | Nr3c1 | | | | | | | | |
| Egr1 | Nrip1 | | | | | | | | |
| F13a1 | Odc1 | | | | | | | | |
| F5 | P2ry10 | | | | | | | | |
| Fcer1g | Paxbp1 | | | | | | | | |
| Fcnb | Pde4d | | | | | | | | |
| Fes | Pja1 | | | | | | | | |
| Fgd4 | Plet1 | | | | | | | | |
| Fpr1 | Polr2a | | | | | | | | |
| Fpr2 | Ppp1r16b | | | | | | | | |
| Fth1 | Prrc2c | | | | | | | | |
| Ftl1 | Ptpn22 | | | | | | | | |
| Gas7 | Rab8b | | | | | | | | |
| Gda | Ramp3 | | | | | | | | |
| Gpr35 | Rel | | | | | | | | |
| Grina | Rnf125 | | | | | | | | |
| Grn | S1pr1 | | | | | | | | |
| Gsn | Samsn1 | | | | | | | | |
| Gsr | Satb1 | | | | | | | | |
| H2-Aa | Scgb1a1 | | | | | | | | |
| Hck | Slc3a2 | | | | | | | | |
| Hdc | Smad3 | | | | | | | | |
| Hp | Snx18 | | | | | | | | |
| Ifitm1 | Socs3 | | | | | | | | |
| Igsf6 | Sqstm1 | | | | | | | | |
| Il13ra1 | Stat3 | | | | | | | | |
| Il1rn | Stat5b | | | | | | | | |
| Il36g | Stk17b | | | | | | | | |
| Itgam | Stk4 | | | | | | | | |
| Itgax | Tbc1d30 | | | | | | | | |
| Jaml | Tex2 | | | | | | | | |
| Kcnj2 | Tgfbr2 | | | | | | | | |
| Klra2 | Thap12 | | | | | | | | |

Lgals3
Lilra6
Litaf
Lpcat2
Lrp1
Lrrc25
Ly6c2
Ly6g
Lyn
Lyst
Marchf1
Mefv
Mgst1
Mmp9
Mpeg1
Myo1f
Nadk
Ncf1
Ncf2
Nfe2
Nfe2l2
Nlrp3
Olfm4
Padi4
Pax5
Pbx1
Pgd
Pglyrp1
Pi16
Pid1
Pilra
Pirb
Pla2g7
Plac8
Plaur
Plbd1
Plcg2
Plek
Plxnb2
Prdx5
Prkar2b
Prkcd
Prss34
Psap
Ptgs2
Ptprj
Ptpro
Pygl
Rassf3
Rassf4
Retnlg
Rnase6
Scd1
Sgms2
Sirpa
Slc11a1
Slc16a3
Slc2a6
Slc4a1
Slpi
Sord
Sorl1
Spi1
Spta1
Sptb
Stom
Syk
Taldo1
Tbc1d8
Thbs1
Themis2
Tifa
Tifab
Tlr2
Tm6sf1
Tmcc1
Tnfaip2
Tnfrsf21
Tpd52
Trem1
Trem3
Trf
Trim30b
Tyrobp
Vwf
Wdfy3
Wfdc21
Xdh
Zeb2
Thrap3
Tinf2
Tmem64
Tob2
Trib2
Trp53inp1
Ubald2
Zc3hav1
Zdhhc23
Zeb1
Zfp281
