## Supplementary Table S2 for "Low-self-reactive, circulation-biased blood regulatory T cells sense danger-associated nucleotides to restrain atherosclerosis"

| Supplementary Table S2, CD38-quartile DEGs in human PB Tregs | | | | | | | | | | | | | | | | | | | | | | | |
|---|---|---|---|---|---|---|---|---|---|---|---|---|---|---|---|---|---|---|---|---|---|---|---|
| Fraction 1–enriched genes | | | Fraction 1–downregulated genes | | | Fraction 2–enriched genes | | | Fraction 2–downregulated genes | | | Fraction 3–enriched genes | | | Fraction 3–downregulated genes | | | Fraction 4–enriched genes | | | Fraction 4–downregulated genes | | |
| gene_symbol | min_padj | min_abs_lfc | gene_symbol | min_padj | min_abs_lfc | gene_symbol | min_padj | min_abs_lfc | gene_symbo | min_padj | min_abs_lfc | gene_symbo | min_padj | min_abs_lfc | gene_symbo | min_padj | min_abs_lfc | gene_symbo | min_padj | min_abs_lfc | gene_symbo | min_padj | min_abs_lfc |
| MIAT | 3.69E-29 | 0.49464273 | GNLY | 1.52E-14 | 0.21465884 | RBM33-DT | 0.01201811 | 1.51542208 | GNLY | 3.49E-10 | 0.21465884 | IL7R | 1.09E-11 | 0.21690046 | ZNF267 | 0.00523257 | 0.39586894 | GNLY | 1.52E-14 | 1.47865374 | TXNIP | 7.29E-44 | 0.51775348 |
| GBP5 | 1.80E-20 | 0.36787584 | SMN1 | 0.00082249 | 0.63537938 | PARD3B | 0.02610727 | 1.2192073 | CD74 | 7.86E-09 | 0.60014538 | ACAP1 | 2.28E-07 | 0.18327777 | SPTLC3 | 0.00597935 | 0.46469627 | FCGR3A | 2.42E-08 | 2.6607443 | CXCR4 | 1.09E-33 | 0.7197867 |
| TMC8 | 1.88E-13 | 0.26435155 | LAMA2 | 0.0027018 | 1.41146941 | FPGT-TNNI3K | 0.02947755 | 1.7083414 | FCGR3A | 2.42E-08 | 2.33334793 | KLF2 | 3.92E-07 | 0.15490652 | NANOS1 | 0.01440614 | 0.88152901 | LILRB2 | 1.53E-07 | 2.3670893 | MIAT | 3.69E-29 | 0.53050111 |
| TIGIT | 9.50E-13 | 0.48273304 | CCR7 | 0.0051192 | 0.22810246 | SPTLC3 | 0.03495662 | 0.19666636 | MYO1F | 2.67E-06 | 0.6639864 | IL16 | 1.41E-06 | 0.03371011 | | | | SRGN | 3.33E-07 | 0.60740577 | BIRC3 | 5.22E-25 | 0.81298396 |
| RESF1 | 2.06E-10 | 0.24569245 | DNALI1 | 0.01011555 | 2.011648 | | | | IL16 | 0.00021662 | 0.16249741 | GNAQ | 5.04E-05 | 0.45204536 | | | | FCN1 | 7.66E-06 | 3.14476317 | GBP5 | 1.80E-20 | 0.36837788 |
| GDPD5 | 8.17E-10 | 0.48499395 | SPTLC3 | 0.02369805 | 0.46469627 | | | | LINC00861 | 0.00264854 | 0.64428923 | GNLY | 0.00015045 | 1.47865374 | | | | CCR9 | 1.61E-05 | 2.73880759 | JUNB | 5.34E-20 | 0.39697203 |
| KDM5D | 8.24E-08 | 0.39322246 | IGHM | 0.04153338 | 0.08806107 | | | | AQP3 | 0.00726169 | 0.55147691 | PIGN | 0.00524688 | 0.2054527 | | | | NKG7 | 1.76E-05 | 2.05207192 | TRAF1 | 6.56E-16 | 0.41047217 |
| NLRC5 | 1.70E-07 | 0.33456631 | PLCD3 | 0.06682164 | 0.15964221 | | | | KLRD1 | 0.00740567 | 0.28537385 | CHRM3-AS2 | 0.00566243 | 0.85100797 | | | | COL5A3 | 5.54E-05 | 1.0020225 | SORL1 | 8.63E-16 | 0.66140962 |
| IL16 | 2.22E-07 | 0.03371011 | | | | | | | AKR1C1 | 0.00890434 | 1.59211888 | HBB | 0.00711682 | 0.35544522 | | | | CCL5 | 8.01E-05 | 1.00228992 | TMC8 | 1.88E-13 | 0.53025552 |
| PLCH2 | 5.72E-07 | 0.7133204 | | | | | | | TC2N | 0.01020765 | 0.44093707 | C11orf21 | 0.01682604 | 0.12368147 | | | | GZMB | 9.75E-05 | 3.07500857 | IL7R | 1.09E-11 | 0.39897968 |
| CYTH4 | 4.58E-06 | 0.42397235 | | | | | | | C11orf21 | 0.01287552 | 0.07558451 | IGHM | 0.0458476 | 0.26953759 | | | | CTSS | 0.0001956 | 0.54358962 | PI16 | 2.22E-11 | 2.03166303 |
| JAML | 1.43E-05 | 0.30804028 | | | | | | | ITGB2 | 0.01450033 | 0.50708807 | | | | | | | MS4A7 | 0.00029611 | 2.63975472 | RESF1 | 2.06E-10 | 0.25242323 |
| SP100 | 0.0001369 | 0.58517742 | | | | | | | ARHGAP9 | 0.0256198 | 0.2772911 | | | | | | | CYBB | 0.00073049 | 2.36931021 | FCRL3 | 5.01E-10 | 0.78121646 |
| F11R | 0.00015045 | 0.69291665 | | | | | | | GIMAP4 | 0.06142029 | 0.41127687 | | | | | | | LST1 | 0.00169847 | 2.26755023 | GCNT4 | 5.01E-10 | 1.39202616 |
| PRICKLE1 | 0.00018805 | 0.19448256 | | | | | | | HBB | 0.06385509 | 0.80953184 | | | | | | | FGR | 0.00187641 | 1.47723275 | GDPD5 | 8.17E-10 | 0.29390802 |
| KCNN4 | 0.00037782 | 0.85175807 | | | | | | | ANKRD44 | 0.06469736 | 0.20337154 | | | | | | | MPEG1 | 0.00197934 | 2.77011328 | PIK3IP1 | 6.19E-09 | 0.3646645 |
| NLRC3 | 0.00145399 | 0.00674596 | | | | | | | | | | | | | | | | SIGLEC10 | 0.00200073 | 2.24542107 | BCL2 | 6.19E-09 | 0.4673445 |
| C11orf21 | 0.00180767 | 0.12368147 | | | | | | | | | | | | | | | | SERPINA1 | 0.00219064 | 2.69205382 | ITK | 1.31E-08 | 0.49464649 |
| IKZF5 | 0.00566736 | 0.61037158 | | | | | | | | | | | | | | | | LILRA1 | 0.00517726 | 3.60022713 | FOXP3 | 1.61E-08 | 0.4676222 |
| STON2 | 0.01021418 | 0.13508449 | | | | | | | | | | | | | | | | SPTLC3 | 0.00597935 | 0.19666636 | KDM5D | 8.24E-08 | 0.408826 |
| HBB | 0.01505294 | 0.35544522 | | | | | | | | | | | | | | | | KLRD1 | 0.00740567 | 0.166596 | NLRC5 | 1.70E-07 | 0.36585553 |
| PIK3R5 | 0.01653033 | 0.08053731 | | | | | | | | | | | | | | | | PRF1 | 0.0138056 | 0.70853636 | IL16 | 2.22E-07 | 0.16249741 |
| RASGRP1 | 0.02012865 | 0.30897125 | | | | | | | | | | | | | | | | CD300E | 0.01505294 | 2.6922813 | ACAP1 | 2.28E-07 | 0.10031879 |
| RBMS2 | 0.04046275 | 0.99467691 | | | | | | | | | | | | | | | | LILRB1 | 0.01584224 | 1.68308248 | KLF2 | 3.92E-07 | 0.33847372 |
| SNAP23 | 0.05225804 | 0.83358294 | | | | | | | | | | | | | | | | NCF2 | 0.01747893 | 2.09096148 | LCP2 | 4.18E-07 | 0.34003928 |
| CRYBG1 | 0.06707398 | 0.76976574 | | | | | | | | | | | | | | | | MPP3 | 0.02478882 | 0.1546176 | LTB | 4.24E-07 | 0.34969757 |
| | | | | | | | | | | | | | | | | | | ITGA4 | 0.03093102 | 0.60363368 | ARHGAP4 | 7.13E-07 | 0.31381938 |
| | | | | | | | | | | | | | | | | | | PECAM1 | 0.03235075 | 0.52783446 | MAP3K1 | 1.01E-06 | 0.36861869 |
| | | | | | | | | | | | | | | | | | | FGFBP2 | 0.03939285 | 1.47044295 | P2RY8 | 3.15E-06 | 0.35894089 |
| | | | | | | | | | | | | | | | | | | | | | CYTH4 | 4.58E-06 | 0.42052522 |
| | | | | | | | | | | | | | | | | | | | | | CD37 | 5.80E-06 | 0.43562561 |
| | | | | | | | | | | | | | | | | | | | | | ADAM8 | 1.12E-05 | 0.62140012 |
| | | | | | | | | | | | | | | | | | | | | | CBX7 | 1.13E-05 | 0.67219826 |
| | | | | | | | | | | | | | | | | | | | | | BCL11B | 3.27E-05 | 0.32875537 |
| | | | | | | | | | | | | | | | | | | | | | GVINP1 | 0.00012076 | 0.42423595 |
| | | | | | | | | | | | | | | | | | | | | | ZAP70 | 0.00018805 | 0.25492045 |
| | | | | | | | | | | | | | | | | | | | | | PRICKLE1 | 0.00018805 | 0.33054416 |
| | | | | | | | | | | | | | | | | | | | | | CRIP2 | 0.00029198 | 0.95490878 |
| | | | | | | | | | | | | | | | | | | | | | PAG1 | 0.00029423 | 0.30397573 |
| | | | | | | | | | | | | | | | | | | | | | TTN | 0.00029601 | 0.60358379 |
| | | | | | | | | | | | | | | | | | | | | | ZMAT1 | 0.00029777 | 0.79776972 |
| | | | | | | | | | | | | | | | | | | | | | MIR3142HG | 0.00106606 | 1.10262835 |
| | | | | | | | | | | | | | | | | | | | | | CSGALNACT | 0.00129022 | 0.65316011 |
| | | | | | | | | | | | | | | | | | | | | | NLRC3 | 0.00145399 | 0.08385806 |
| | | | | | | | | | | | | | | | | | | | | | TAGAP | 0.00163491 | 0.37801519 |
| | | | | | | | | | | | | | | | | | | | | | C11orf21 | 0.00180767 | 0.07558451 |
| | | | | | | | | | | | | | | | | | | | | | CISH | 0.00180767 | 0.70469884 |
| | | | | | | | | | | | | | | | | | | | | | UBASH3A | 0.00321783 | 0.55645719 |
| | | | | | | | | | | | | | | | | | | | | | VWA7 | 0.00447006 | 3.11912829 |
| | | | | | | | | | | | | | | | | | | | | | NA | 0.00492912 | 1.43492106 |
| | | | | | | | | | | | | | | | | | | | | | IL11RA | 0.00640852 | 0.1679972 |
| | | | | | | | | | | | | | | | | | | | | | HBB | 0.00711682 | 0.80953184 |
| | | | | | | | | | | | | | | | | | | | | | SIDT1 | 0.00811682 | 1.32927266 |
| | | | | | | | | | | | | | | | | | | | | | JMJD7-PLA2 | 0.01517587 | 0.38477789 |
| | | | | | | | | | | | | | | | | | | | | | IFNAR1 | 0.01673614 | 0.50784395 |
| | | | | | | | | | | | | | | | | | | | | | CRBN | 0.01789202 | 0.43580907 |
| | | | | | | | | | | | | | | | | | | | | | KCNA3 | 0.03704282 | 0.52006189 |
| | | | | | | | | | | | | | | | | | | | | | RAB8B | 0.04286124 | 0.33741276 |
| | | | | | | | | | | | | | | | | | | | | | SELP | 0.08629497 | 1.80663355 |
| | | | | | | | | | | | | | | | | | | | | | SPATA7 | 0.08968326 | 0.61630319 |
